# OcuPair, a novel photo-crosslinkable dendrimer-hyaluronic acid hydrogel bandage/bioadhesive for corneal injuries and temporary corneal repair

**DOI:** 10.1101/2025.01.08.632042

**Authors:** Siva P. Kambhampati, Rishi Sharma, Hui Lin, Santiago Appiani, Jeffrey L. Cleland, Samuel C Yiu, Rangaramanujam M. Kannan

**Affiliations:** Center for Nanomedicine, Wilmer Eye Institute, Department of Ophthalmology, Johns Hopkins University School of Medicine, Baltimore, MD, USA; Ashvattha Therapeutics, Inc., Baltimore, MD, USA

**Keywords:** Open-globe battlefield corneal injuries, bioadhesive hydrogels, PAMAM dendrimer, hyaluronic acid, methacrylated dendrimer, methacrylate hyaluronic acid, photo-crosslinkable hydrogels, biopolymers, hydrogel bandage

## Abstract

Traumatic corneal injuries are a leading cause of blindness among military personnel. These injuries need immediate attention at the combat zone, but treatment options are limited as life-saving measures are often prioritized. To address this critical gap, we have developed Ocupair^TM^, a two-component hydrogel system that consists of (i) an injectable viscoelastic filler that stabilizes the ocular cavity and prevents hypotony. (ii) An *in-situ* photo-curable adhesive hydrogel comprising of methacrylated PAMAM dendrimer and hyaluronic acid engineered to form a transparent, flexible and robust bandage within 90 seconds, adhering to corneal surface and ensuring a water-tight seal securing full-thickness corneal wounds. *Ex vivo* studies demonstrated that the adhesive hydrogel is mechanically robust and withstands intraocular pressures beyond physiological range. In a rabbit corneal injury model, OcuPair™ effectively seals complex full thickness wounds and preserves the eye with favorable clinical outcomes for 5 days with no toxicity over 30 days. In this study, we have validated the pilot scale synthesis, formulation optimization, GMP scale-up and IDE-enabling GLP toxicity, essential for clinical translation as a battlefield-ready solution.

## 1. Introduction

Battlefield related traumatic eye injuries (TEI) are one of the leading causes of blindness in military personnel [1, 2]. It is estimated that more than 20% of the US soldiers evacuated from Operation Iraqi Freedom/Operation Enduring Freedom had significant open globe injuries and this number is estimated to increase in coming years [3, 4]. Despite the use of ocular protection, improvements in the development of Improvised Explosive Devices (IEDs) have increased the risk and severity of blast/traumatic injuries on the battlefield. [5, 6]. These type of open globe injuries involve high morbidity with complex multiple full-thickness lacerations in the cornea, severe ocular hypotony and iris prolapse, and loss of intraocular fluids due to gaping corneal wounds, which results in damage to the posterior segment, including retinal detachment and choroidal hemorrhage [3, 7, 8]. Although less frequent, such scenarios can also occur in civilian trauma and mass casualty events [4, 9]. Much of the life-saving efforts by forward surgical team in the combat theater of operations are focused on treating other injuries that involve the head, extremities, and trunk [10]. In such cases, ocular trauma may be often overlooked, resulting in delayed treatment [11]. Another important problem is that such injuries involve pathological events such as acute inflammation and infections, which trigger downstream ocular tissue hemostasis, resulting in poor visual outcomes or permanent vision loss [9].

Current options for addressing, treating, and managing TEIs involve sutures and mechanical barrier therapies such as pressure patching or adhesives [12]. Sutures require additional microsurgical instrumentation accompanied by surgical microscope which may not be a viable option in role of care (ROC) near battlefield combat zones. Adhesives such as fibrin glue and cyanoacrylates have been explored but are limited to small corneal aberrations, surgical incisions and small corneal perforations. They may not be suitable for full thickness injures and globe injuries with missing tissues [9, 13]. Additionally, for cyanoacrylate glue, cytotoxicity and chronic inflammation induced by the degraded by products are a notable drawback [14, 15]. Since the eye is a highly sensitive and delicate organ, the use of cyanoacrylates to address small corneal perforations is somewhat challenging. Accidental deposition of the product to intraocular compartments can result in serious toxic effects [16–18]. Therefore, careful selection of materials compatible with ocular tissues to develop adhesives for corneal applications is highly crucial. To date, there are no U.S. FDA approved solutions for using medical adhesives for closure of full thickness corneal and corneo-scleral injuries. Recent studies by multiple research groups have explored photo-crosslinkable hydrogels for ocular wound closures, that have shown promise [19–21]. There is a need for interventions that can temporarily stabilize the eye and secure corneal wounds to protect the eye for 24-72 hours until appropriate clinical treatments or operative procedures can be performed once the injured army/civilian personnel have been transported to higher role of care (ROC) facilities. Further, it will be highly beneficial if new solutions can address the following: (i) easily applicable in the ROC setting near the combat zone without the requirement for sophisticated instrumentation, (ii) creates a water-tight seal securing full thickness wounds and withstands increasing intraocular pressure (IOP), (iii) be transparent so that the intraocular structures and damage can be assessed, (iv) can be easily removed by surgical instruments in higher ROC settings, and (v) that can address globe hypotony if required.

We have developed OcuPair^TM^, a two-component system consisting of (i) an injectable filler consisting of a hyaluronic acid viscoelastic that can be injected into the ocular cavity to address globe hypotony and to contain the intraocular structures and prevent iris prolapse, (ii) adhesive hydrogel, a viscous fluid that is a mixture of methacrylate-functionalized hyaluronic acid (HA-MA) and a methacrylate-functionalized hydroxyl polyamidoamine dendrimer (D-MA) which can be easily applied over the cornea and photocured *in situ* to form a transparent hydrogel bandage over the cornea. This system leverages the unique properties of a linear biopolymer hyaluronic acid (HA) and a branched globular hydroxyl PAMAM dendrimer. HA is an integral component of vitreous humor, aqueous fluid, and the corneal stroma [22, 23]. HA also influences several cellular mechanisms for wound healing by recruiting fibroblasts and promoting re-epithelization at the wound site, and aiding the production of extracellular matrix for accelerating stromal expansion [24, 25]. It has also been reported to have anti-inflammatory and antioxidant properties that contributes to cell proliferation [26–28]. HA also has unique viscoelastic properties with high water retention capacity contributing to its shear thinning, hydration and lubricating properties. Hence, HA is being widely used in various ocular formulations such as injectable viscoelastic gels and eyedrops [29, 30]. On the other hand, Generation 4 PAMAM hydroxyl dendrimers (D-OH) are well defined, tree-like branched globular nanostructured polymers with highly tunable surface chemistry. D-OH dendrimers are well tolerated and have favorable non-toxic profile and have been explored at depth by our group for ocular applications demonstrating selective targeting, localization and precisely delivering therapeutic drugs and biologics into reactive/activated microglia, macrophages and hypertrophic retinal pigment epithelium without any targeting in multiple small and large animal models. They have shown promise in early Phase 2 trials [31–41].

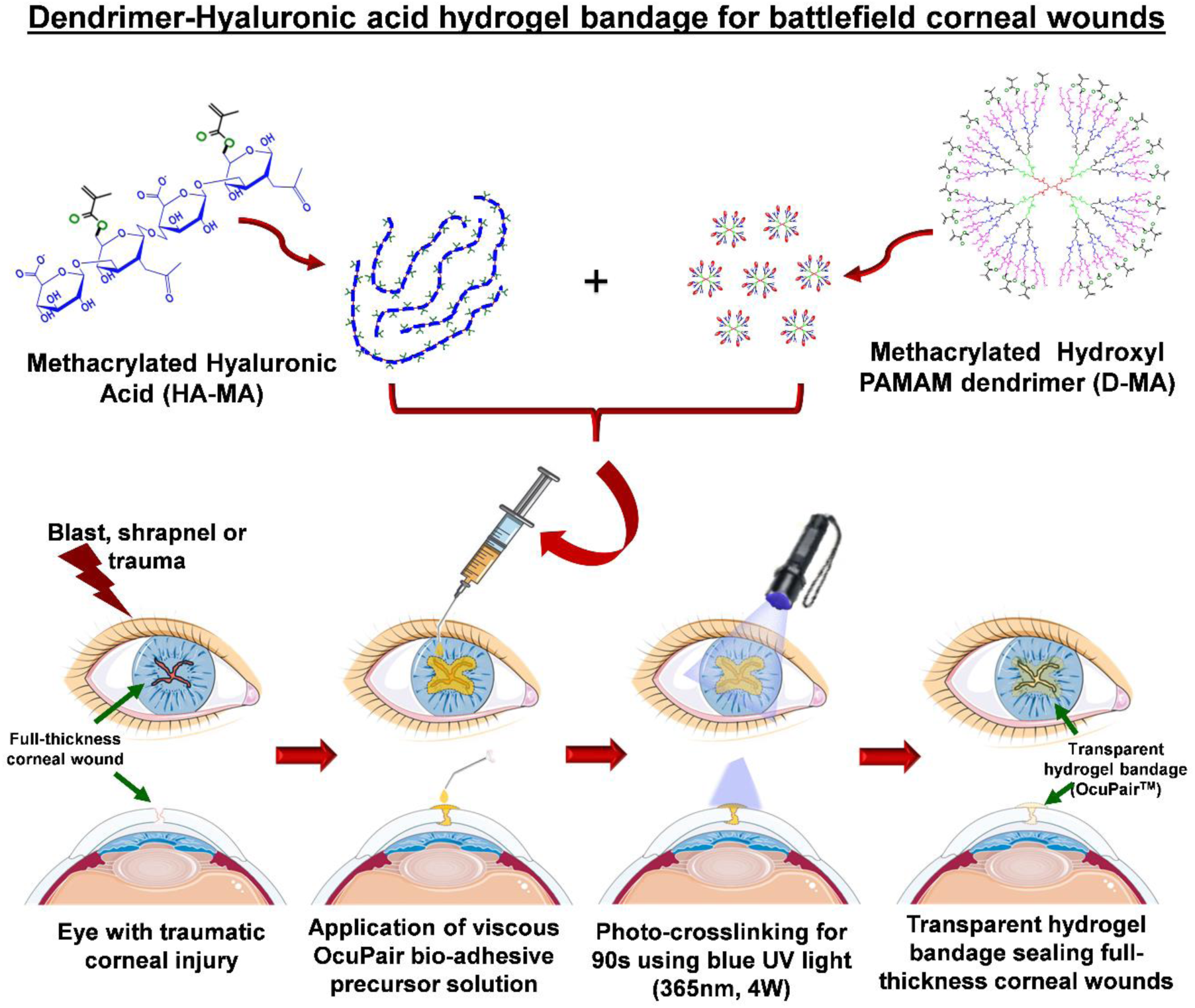
Table of Contents/Schematic. A schematic diagram showing the OcuPair adhesive hydrogel formulation loaded into final delivery device and applied over the full-thickness corneal wound in an in vivo rabbit model of corneal injury followed by in situ crosslinking to form a transparent hydrogel bandage sealing the wound. Parts of the figure were drawn by using pictures from Servier Medical Art (http://smart.servier.com/), licensed under a Creative Commons Attribution 4.0 Unported License (https://creativecommons.org/licenses/by/4.0/).

In this study, we developed an adhesive hydrogel (OcuPair adhesive hydrogel) by harnessing the branched architecture and multivalent nature of dendrimers which enables multiple copies of photo-crosslinkable groups on to the dendrimer surface enabling high crosslinking density enabling tailorable mechanical strength to the hydrogel, dictated by the physician. Conversely, hyaluronic acid possesses mucoadhesive and viscoelastic properties with multiple side hydroxyl groups amenable for methacrylate functionalization contributing flexibility and adhesiveness to the hydrogel. We report the synthesis and physiochemical characterization of D-MA and HA-MA amenable for scale-up synthesis, formulation development and material characterization of adhesive hydrogel. Feasibility studies of OcuPair performance were evaluated using *ex vivo* and *in vivo* models of corneal traumatic injury in rabbits, assessed against clinically relevant benchmark.

## 2. Results and Discussions

### 2.1. Synthesis and characterization of D-MA and HA-MA conjugates

Dendrimer-Methacrylate (D-MA) conjugates were synthesized using single step coupling reactions between surface hydroxyl groups (-OH) of PAMAM dendrimers and methacrylic anhydride in the presence of DMAP under nitrogen atmosphere (**Figure 1A**). The product was analyzed using ^1^H NMR technique (**Figure 1B**) to confirm the structure and target loading of methacrylate on the dendrimer. In the **^1^H NMR spectrum**, once the methacrylic groups are covalently attached on to the surface of dendrimer, a new –CH_2_ peak appears at δ 4.1 ppm. This peak corresponds to –CH_2_ groups present on dendrimer which participates in the formation of ester linkage with methacrylate group (**Figure 1B**). The methyl (-CH_3_) of the methacrylic group appears at δ 1.9 ppm. The methylene protons (-CH_2_) of methacrylic group gives two broad singlets at δ 6.0 ppm and δ 5.6 ppm respectively (**Figure 1B**). The degree of substitution of the methacrylic group was calculated by using the proton integration technique. By integrating of these signature peaks of dendrimer (internal amide) and methacrylic group (CH_3_ & CH_2_), we estimated that ∼16-20 molecules of methacrylate were conjugated to one dendrimer molecule (**Figure 1B**). By following the above synthesis procedure, we have synthesized multiple pilot-scale batches of D-MA demonstrating consistent degree of methacrylation and purity (**Figure S1, supporting information**). **HPLC** analysis of D-MA demonstrated a broad peak at 12.9 min which is different from starting dendrimer at 8 min (data not shown) with a purity of >95% (**Figure S2, supporting information**). The conjugates were readily soluble in water, PBS buffer and saline.

**Figure 1:**
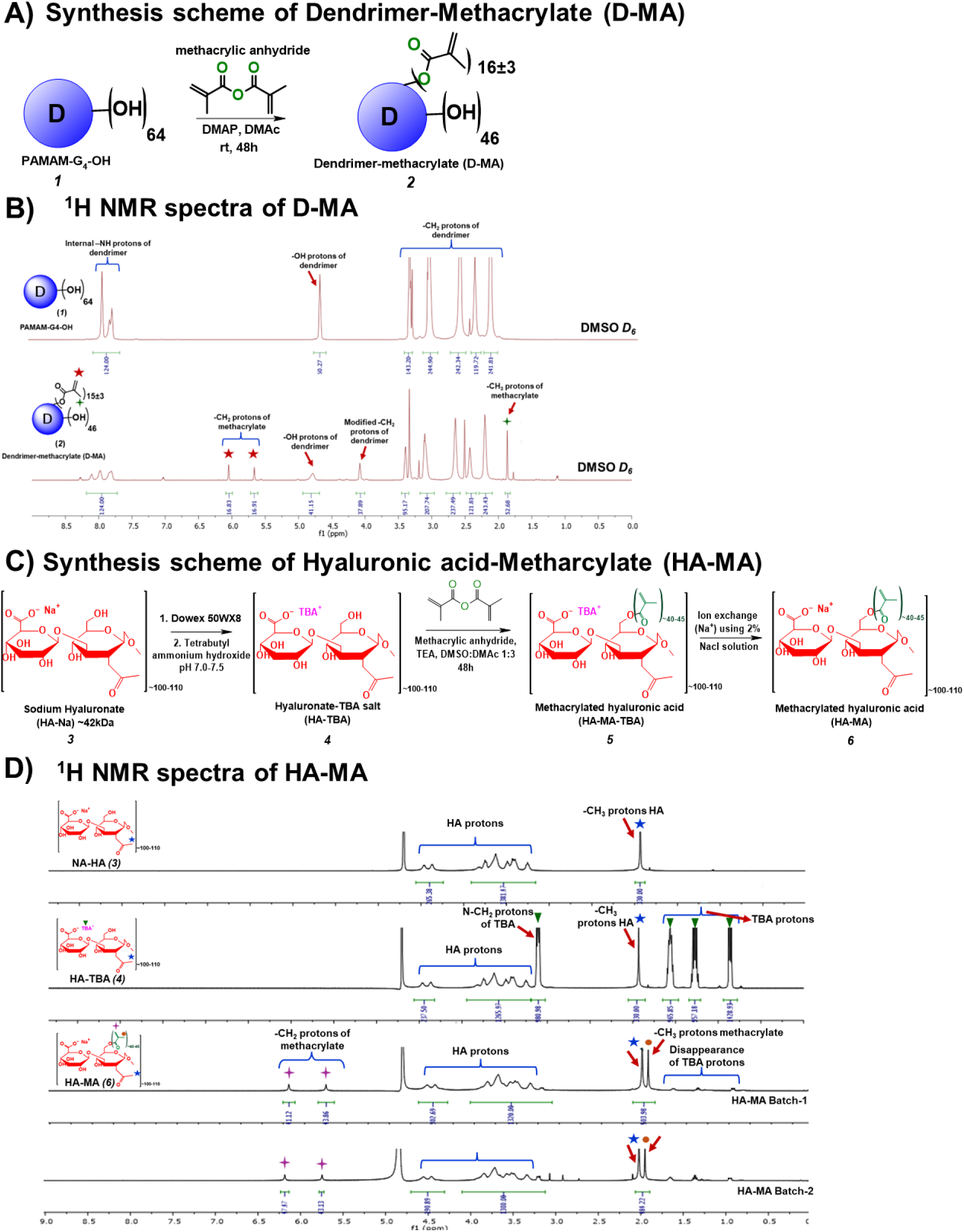
Synthesis and NMR characterization for OcuPair adhesive hydrogel components (D-MA and HA-MA). **A)** Synthesis scheme of Dendrimer-Methacrylate (D-MA) conjugate. **B)** 1H NMR spectrum of G4-OH dendrimer and D-MA. **C)** Synthesis scheme of Hyaluronic Acid-Methacrylate (HA-MA) conjugate. **D)** 1H NMR spectrum of Na-HA, HA-TBA and HA-MA (2 consecutive batches) conjugates.

Methacrylated hyaluronic acid synthesis has been extensively investigated by several research groups, typically conducted in aqueous solutions or water/organic solvent blends [43–45]. However, these reported procedures often (i) yield inconsistent results with relatively low degrees of substitution and (ii) may require adding higher mole equivalents for methacrylate and longer reaction duration. To have a consistent and high degree of substitution of methacrylate groups to the –OH groups of hyaluronic acid, we adopted an alternative synthesis route by modifying a previously reported procedure [37, 42] (**Figure 1C**). In the first step, the commercially available sodium salt of hyaluronic acid (∼42 kDa) (purchased from Lifecore Biomedical) is converted to tetrabutylammonium salt (HA-TBA) using an ion exchange method. This conversion transforms the water-soluble sodium salt of HA into HA-TBA salt, which is soluble in organic solvents, thus facilitating further chemical modifications in inert atmosphere and anhydrous conditions. To prepare HA-TBA, a 2% solution of sodium hyaluronate (Na-HA) in deionized water is treated with Dowex 50W proton exchange resin (strongly acidic) for 5 hours. After the exchange, the resin is filtered out, and the filtrate is titrated to a pH of 7.1-7.5 with a 1.0 M solution of tetrabutylammonium hydroxide in methanol. The resulting solution is then frozen at –80°C and lyophilized. The exchange of Na+ for TBA+ ions is confirmed by ¹H NMR analysis, which shows new peaks corresponding to tetrabutylammonium: a triplet at δ 0.91 ppm (CH₃), a multiplet at δ 1.38-1.35 ppm (CH₂), a multiplet at δ 1.66-1.63 ppm (CH₂), and a triplet at δ 3.21-3.18 ppm (CH₂) (**Figure 1D**).

In the second step, methacrylate groups are introduced to HA-TBA to obtain MA-HA-TBA. We use a highly efficient and milder anhydride chemistry for the partial esterification of HA-TBA. The target conjugation is approximately 40±5 methacrylate molecules per dendrimer. HA-TBA was dissolved in a DMSO/DMAc mixture and reacted with methacrylic anhydride in the presence of triethylamine (TEA) as base. The reaction mixture is stirred in the dark for 48 hours. HA-MA-TBA is then purified using tangential flow filtration (TFF) with a 10 kDa TFF cartridge, and the retentate is collected in DMAc.

In ¹H NMR (**Figure 1D**), appearance of new peaks corresponding to methacrylate appear, for the methyl group of the methacrylic group appears at δ 1.85 ppm. The CH₂ protons of the methacrylic group show two broad singlets at δ 6.0 ppm and δ 5.6 ppm. In the third step, the TBA ions are fully exchanged with Na+ using a 2% NaCl solution followed by dialysis against water using TFF to remove free TBA ions The MA-HA-TBA solution in DMA is replaced with 2% NaCl solution via TFF and the resultant solution was lyophilized. The removal of TBA and the successful conjugation of HA are confirmed by NMR analysis. ^1^H NMR the peaks corresponding to the tetrabutylammonium group (δ 0.91 ppm, δ 1.38-1.35 ppm, δ 1.66-1.63 ppm, and δ 3.21-3.18 ppm) disappear, and the peaks pertaining to methacrylate groups remain. The degree of substitution of the methacrylic group is calculated by integrating these signature peaks, revealing that approximately 40±5 methacrylate groups were introduced with a degree of methacrylation of ∼50%. Using the same procedure, we have synthesized multiple pilot-scale batches of HA-MA with consistent degree of substitution and yields (**Figure S3, supporting information**) demonstrates the robustness and efficiency of the optimized synthesis protocol. Further, the HPLC spectra demonstrates a broad peak at 2.9 min with a purity >97% (**Figure S4, Supporting information**). The optimized synthesis route for D-MA and HA-MA were directly used for the scale-up synthesis by the CMO’s under GMP settings for developing GLP engineering batches.

### 2.2. Challenges associated with the scalable synthesis of D-MA and HA-MA, and the optimized method

Initially, we employed traditional coupling conditions for the esterification of methacrylate groups on the PAMAM dendrimer surface, reacting methacrylic acid with EDC and 4-Dimethylaminopyridine (DMAP). While effective on a 500 mg scale, this method encountered significant issues during scale-up to gram quantities. Specifically, multiple 20-gram batches resulted in the formation of a non-dissolvable white mass due to the premature polymerization of methacrylate groups present on the PAMAM G4-OH dendrimer surface. Methacrylates are generally prone to premature polymerization during reactions, storage, or transportation if kept without stabilizers, leading to handling difficulties and compromised product quality. To resolve this issue, we incorporated Hydroquinone Monomethyl Ether (MEHQ) directly into the reaction mixture, ensured the exclusion of oxygen, and used MEHQ during the TFF purification process. Additionally, the entire synthesis and purification process was conducted in the dark to prevent polymerization. The methacrylate loading was precisely controlled by the amount of anhydride equivalents used and the residual methanol present in the reaction. Another challenge was the incomplete removal of DMAP-related impurities, likely encapsulated within the dendrimer. We addressed this by optimizing the purification process using TFF, with 1 L of each feed solution (DI water, 2 wt% brine, and DI water) for every 2 g of PAMAM dendrimer. Purified D-MA was found to be unstable in deionized water at 2-8 °C. Therefore, it was stored in a low pH phosphate buffer saline (pH: 5.5) to ensure stability.

Similar stability-related challenges were encountered for the scale-up of the HA-MA as well but to a lesser extent. To avoid the pre-mature polymerization of HA-MA during the manufacturing process, we added MEHQ at a concentration of ∼100ppm during the entire purification process. These optimized procedures for D-MA and HA-MA consistently produced high-quality products with the desired loading in a reproducible fashion. The standardized synthesis routes on 20-gram pilot-scale were successfully transferred to a contract research manufacturing company for large kilogram scale production of high-purity D-MA and HA-MA.

### 2.3. Formulation and characterization of injectable filler gel (component 1)

The purpose of the injectable filler gel (component 1) in the OcuPair viscoelastic substitute that can be injected into the aqueous chamber to regain the globe architecture and avoid iris prolapse and hypotony. We have designed and formulated the injectable hydrogel by considering following parameters: (i) low viscosity so that the injectable gel is dispersive, and it easily fills the crevices of the intraocular contents to regain the globe architecture and avoid hypotony; (ii) lower concentration compared to commercially available viscoelastic (e.g. Healon EndoCoat is 3% and ProVisc is 3.5%) so that it integrates well with forming aqueous humor, and (iii) use of low molecular weight hyaluronic acid to avoid trabecular meshwork blockage [46–48]. The 2% (20mg/mL, ∼700 kDa Na-HA in phosphate buffer) solution formed a clear transparent viscous solution (**Figure S5 A, Supporting information**). The shear thinning properties of the injectable filler gel was evaluated using dynamic shear frequency and compared it with the commercially available dispersive viscoelastic Healon^®^ EndoCoat. At the low shear rate of 0.01 Hz, the complex viscosity (zero shear viscosity) of the injectable hydrogel is ∼15,685 ± 110 cps which is significantly lower than Healon® EndoCoat (∼50444 ± 852 cps at 0.01 Hz) (p < 0.01, n=4) (**Figure S5 B and C, Supporting information**). The apparent viscosity at a constant shear rate of 100 s^-1^ for the injectable filler gel using capillary viscometer also demonstrated similar results with viscosity of ∼1742 ± 3.2 cps for injectable filler gel whereas the viscosity of the than Healon® EndoCoat was ∼5245 ± 34 cps (p < 0.01, n = 3) (**Figure S5 D, Supporting information**). Sterilization of the injectable filler gel did not alter the viscosity of the injectable filler gel (∼1847 ± 5.2 cps at 100 s^-1^). When loaded into the syringe equipped with 30G cannula, the injectable gel was easily injectable with minimal pressure (assessed qualitatively) (**Figure S5 E, supporting information**). The osmolarity of the injectable filler gel is ∼305 ± 7.7 mOsm/kg and the pH were ∼6.8. The injectable gel can be easily loaded into a 3 mL glass syringe can be easily injectable/extruded using a 30G or 27G anterior chamber cannula and comparable to that of Healon® EndoCoat.

### 2.4. Formulation and characterization of adhesive hydrogel (component 2)

The adhesive hydrogel contains three component solutions: (i) methacrylated dendrimer (D-MA), (ii) methacrylated hyaluronic acid (HA-MA), and (iii) Irgacure 2959 photo-initiator (catalytic amount). The adhesive hydrogel formulation was optimized by using parameters such as viscosity, gelation time, mechanical strength, and flexibility. Both D-MA and HA-MA were readily soluble in phosphate buffer at a concentration of 300 mg/mL and 190 mg/mL respectively. The viscosity of the individual solutions was assessed using capillary micro-viscometry at constant shear rate of 100 s^-1^. The apparent viscosity of D-MA solution at 300 mg/mL was ∼96.8 ± 6.6 cps and for HA-MA solution at concentration of 190 mg/mL was ∼7306.5 ± 336.5 cps **(Figure S5 F, Supporting information)**. Different combinations of mixing ratios D-MA: HA-MA such as 70:30, 50:50, and 30:70 was evaluated in the process of formulation optimization. The 70:30 and 50:50 mixing ratios gelled within 45 seconds but yielded brittle hydrogels and can be attributed to the higher D-MA concentration which contributes to higher crosslinking density (**Table S1, Supporting information**). The 30:70 mixing ratio resulted in solution of viscosity of ∼5205.8 ± 190 cps (**Figure S5 F, Supporting information**). This viscosity of the solution was appropriate, and once applied over the cornea the solution stayed over the incision area on the cornea for ∼15-20 seconds (assessed qualitatively using visual inspection and timer) thereby providing time for photo-crosslinking enabled light radiation by the attending personnel. The formation of adhesive hydrogel and gelation kinetics were assessed using dynamic time sweep rheology at constant strain of 1%. Before blue UV light treatment (until 290 seconds), the 30:70 solution during the liquid phase the adhesive hydrogel solution both storage modulus (G’) and loss modulus (G’’) were low, characteristic of the low solution viscosity before gelling (**Figure 2A**). After UV irradiation, within 20-30 seconds, storage modulus, loss, and complex viscosity increased gradually suggesting the crosslinking reaction. The cross-over of G’ and G” happens within ∼30-45 seconds after UV irradiation and within 90 seconds the storage modulus (G’) increases and the loss modulus stabilizes such that G’>>G” (G’ = ∼35000 ± 350 Pa, G” = ∼1020 ± 92 Pa) (**Figure 2A**), suggesting rapid gelation. The complex viscosity also increased gradually reaching ∼35700 ± 720 Pa.s and stabilizes beyond suggesting the solution is completely crosslinked and confirming gel formation and has viscoelastic properties (**Figure 2A**). The adhesive hydrogel formed with 30:70 mixing ratio was transparent, flexible, transmits light, easily handled using common surgical instruments (**Figure 2B**). Further, a 0.02% fluorescein was included in the formulation to enable identification of the hydrogel bandage over the injured cornea by the ophthalmic surgeon at the operation theater (**Figure 2B**). This formulation once applied over the porcine corneal surface followed by blue UV light irradiation for 90 seconds forms a transparent hydrogel bandage securing and sealing a full thickness corneal incision and can be easily peeled from the corneal surface using common surgical instruments (**Figure 2C and supplementary section Video 3**). The optimized formulation and its process were successfully transferred to a CMO for formulation under sterile settings and filling into final delivery devices. The OcuPair adhesive hydrogel formulation developed by CMO using D-MA and HA-MA GLP batches demonstrated similar viscosities and gelation kinetics suggesting repeatability and robustness of the formulation and its development process (**Figure 2A**).

**Figure 2:**
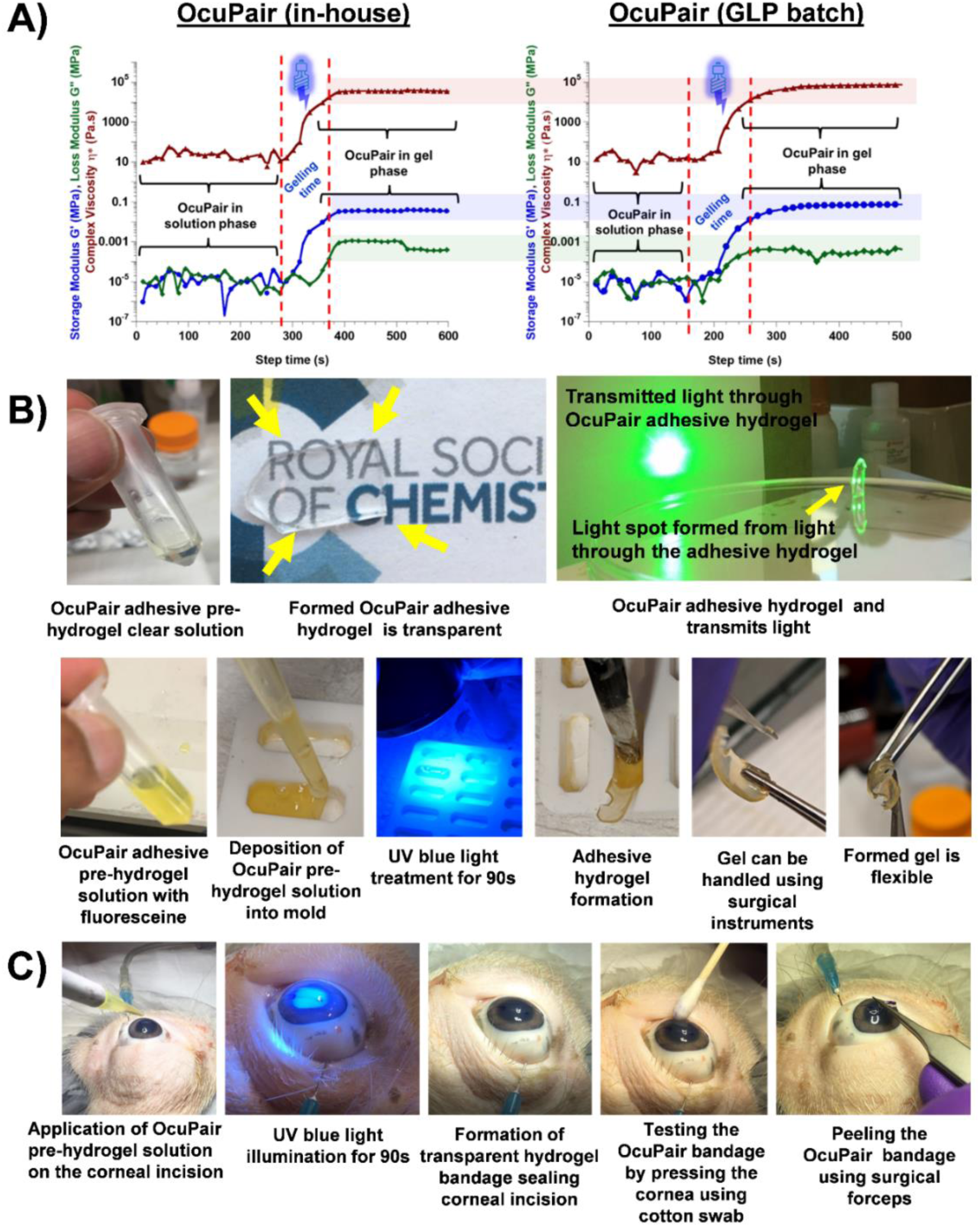
Characterization of OcuPair adhesive hydrogel formulation and gelling kinetics. **A)** Rheological assessment of adhesive hydrogel formation using dynamic time sweep photo-rheology. The pre-polymer viscous solution has low storage modulus (G’) and loss modulus (G”). Upon UV light treatment for 90 seconds, G’ overcomes the G” resulting in a cross-over (happened around 35-45 seconds) followed by steady increase of complex viscosity (η) leading to stabilization of G’, G” and complex viscosity suggestive of completion of gelation. The OcuPair adhesive hydrogel formulation prepared from a GLP batch also demonstrated a similar trend. **B)** A 70:30 mixing ratio of HA-MA and DMA resulted in transparent viscous solution and upon photo-crosslinking for 90 seconds yielded transparent, flexible and sticky hydrogels. **C)** Representative images demonstrating the procedure and steps for sealing corneal incision on *ex vivo* porcine eyes. *In situ* application OcuPair pre-polymer solution over the cornea with a full-thickness linear incision followed by photo-crosslinking resulted in a transparent hydrogel bandage that seals and secures the corneal wound.

### 2.5. Shelf-life stability of D-MA and HA-MA conjugates (4°C, 25°C and 40°C) up to 3 months

We conducted the storage stability of D-MA and HA-MA samples over three months at three different temperatures: 4°C, 25°C, and 40°C in the solution formulation form. A 300 mg/mL and 190 mg/mL of D-MA and HA-MA respectively was prepared in 10mM phosphate buffer solution, aligning with the desired final concentration of the product. The samples were stored in amber-colored vials under argon atmosphere at the desired temperatures. The samples were assessed using HPLC, pH measurements, osmolality, and ^1^H NMR.

D-MA remains relatively stable at 25°C for up to three months, with a slight decrease in HPLC purity from 97.3% to 92.3% by the end of the period. We observed less than 5% methacrylic acid hydrolyzed from the D-MA in HPLC (**Figure S6, supporting information**) and detected in ^1^H NMR (**Figure 3A**). However, significant degradation was observed at 40°C, over 3 months. NMR analysis indicated that the degradation at 40°C is due to the cleavage of methacrylic acid from the D-MA conjugate (**Figure 3A**). Figure 3A shows a decrease in the number of protons corresponding to conjugated methacrylic acid and appearance of new peaks corresponding to cleaved methacrylic acid protons at 6.0, 5.6, and 1.8 ppm (**Figure 3A, red arrows**). Further, decrease in number of protons corresponding to modified methylene (-CH_2_) that are formed due to conjugation of methacrylic acid to the surface hydroxyl groups of dendrimer (**Figure 3A**). These results were corroborated by HPLC, which showed a major impurity, matching the HPLC spectrum of free methacrylic acid (retention time of 7.15 minutes), continuously increasing at 40°C (**Figure S6, Supporting information**). Additionally, no significant changes were observed in the pH of samples at all three temperatures even after 3 months. The pH of the solution at time (t=0h) was 7.59 which changed to 7.31, 7.47, and 7.23 for the samples at 4, 25, and 40°C after 3 months respectively. Finally, the osmolality measurements revealed an increase in osmolality for the samples stored at 25, and 40°C after 3 months, but no significant change was observed for the samples stored at 4°C (**Table S2, Supporting information**). At time (t=0), osmolality of the solution was 201mosm/kg, which changed to 205, 249, and 356 for the samples at 4, 25, and 40°C after 3 months respectively (**Table S2**).

**Figure 3:**
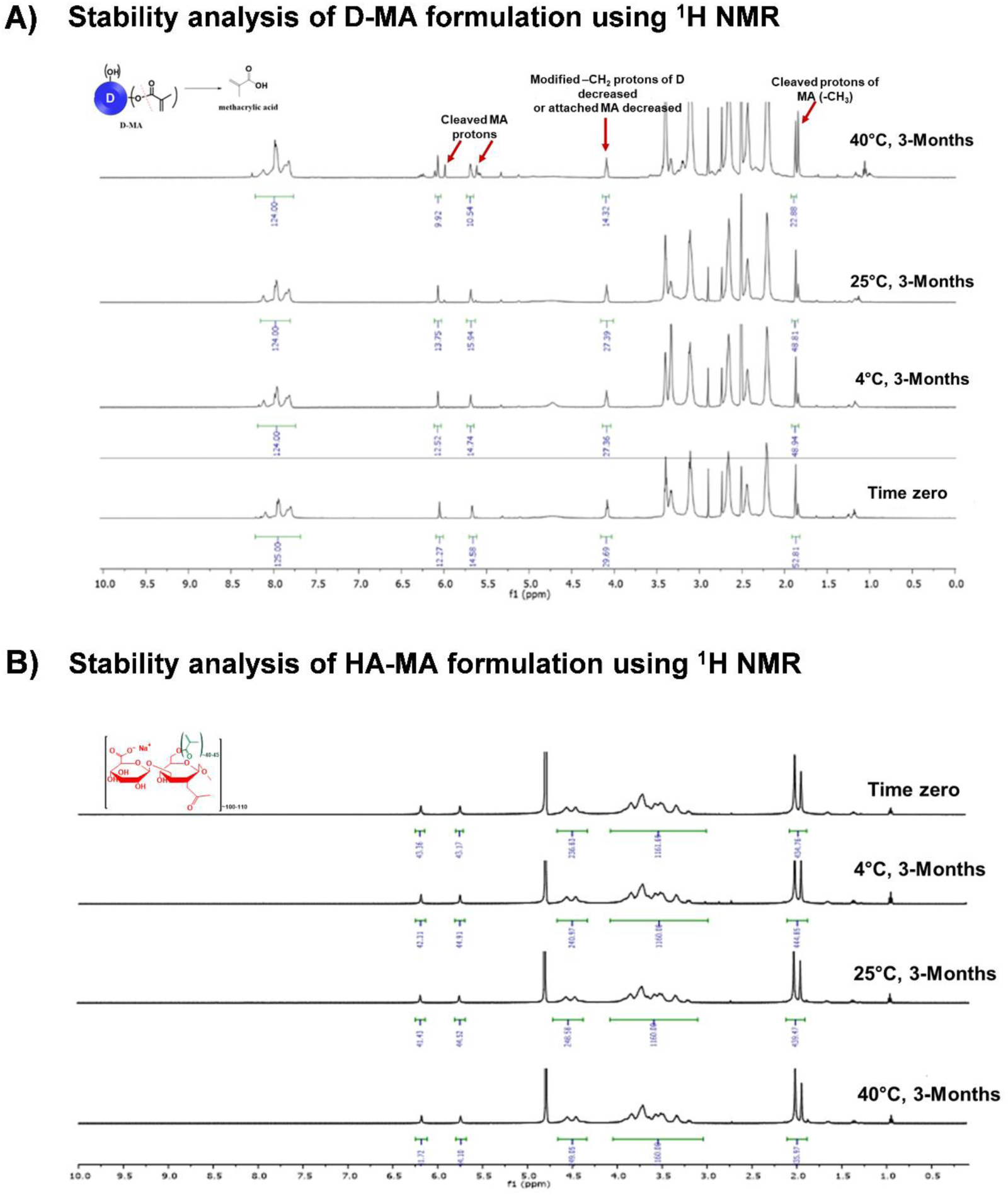
Stability evaluation of OcuPair adhesive hydrogel components using 1H NMR. **A)** 1H NMR spectra of D-MA (300mg/mL) in phosphate buffer (pH 6) stored at 4oC, 25oC, and 40oC at 3-month time point compared with time zero. **B)** 1H NMR spectra of HA-MA solution (190mg/mL) in phosphate buffer (pH 6) stored at 4oC, 25oC, and 40oC at 3-month time point compared with time zero.

Unlike D-MA, HA-MA solution remains highly stable at 4°C, 25°C, and 40°C for up to three months. No significant changes in the structure or the presence of degradation products were observed by ^1^H NMR and HPLC, even at accelerated conditions (40°C) after three months (**Figures 3B and S7 in supporting information**). The methacrylate group, when attached to hyaluronic acid (HA), exhibits greater stability. Unlike D-MA, which we observed was hydrolyzing during synthesis, HA-MA did not undergo polymerization or hydrolysis. We attribute the enhanced stability of methacrylate on hyaluronic acid to the linear structure of hyaluronic acid itself. The methacrylate groups are spaced far apart, minimizing intramolecular interactions between them. Conversely, generation 4 dendrimers are spherical in shape with a diameter of approximately 4nm. Here, methacrylates are closely situated, leading to increased instability and enhanced intramolecular polymerization. Additionally, no significant change was observed in the pH of HA-MA samples at all three temperatures even after 3 months. The pH of the solution at time (t=0h) was 7.6 which changed to 7.4, 7.3, and 7.7 for the samples at 4, 25, and 40°C after 3 months (**Table S2, supplementary information**). The results from the stability studies revealed that the D-MA and HA-MA solution formulations are very stable at 4°C, as indicated by HPLC purity, ^1^H NMR characterization, pH, and osmolality data. Hence, we planned to store the product loaded into final delivery devices at this temperature.

### 2.6. OcuPair adhesive hydrogel seals full thickness corneal wounds and withstand high IOP in porcine and rabbit eyeballs ex vivo

We assessed the burst pressure or sealing capability of OcuPair *ex vivo* using enucleated rabbit and porcine eyes using modified ASTM standard test, F2392-04. Different wound architectures collectively mimicking several aspects of the battlefield injury were created on the porcine and rabbit corneas (**Figure 4A and Table 1**). The adhesive hydrogel pre-solution when applied over the corneal incision stays on the incision site for 15-20 seconds before flowing away. This gives ample time to photo-cure, using blue UV light and within 90 seconds upon light exposure the solution gels form a transparent hydrogel bandage sealing the full thickness wounds immediately (**Figure 4A**). We showed that the two-step process for sealing and stabilizing the incision was effective.

**Figure 4:**
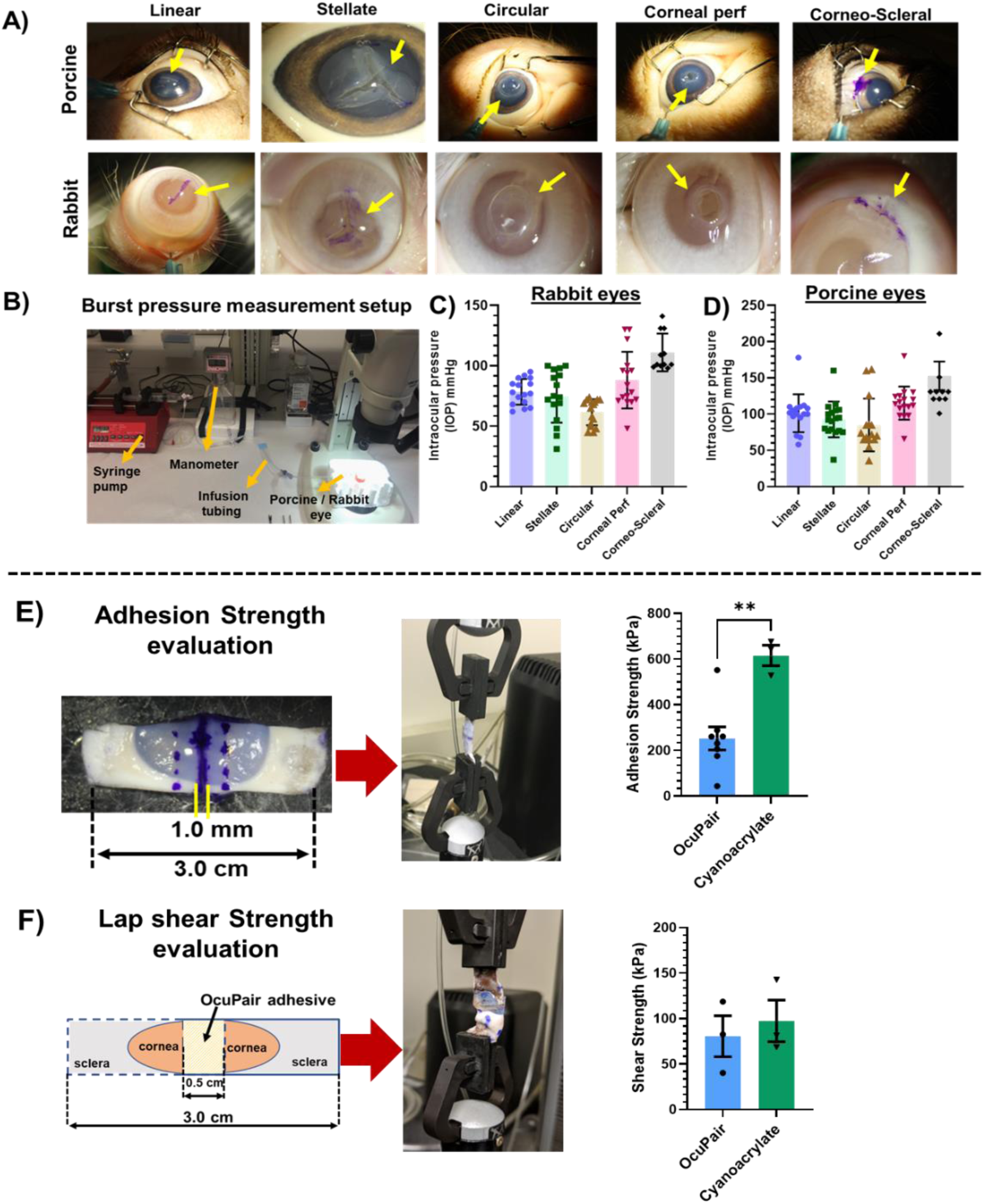
Evaluation of adhesion and mechanical properties in *ex vivo* setting using porcine and rabbit eyes and porcine corneal tissue strips as biological substrates. **A**) Representative images of porcine and rabbit eyeballs with different full-thickness corneal wounds sealed with OcuPair adhesive hydrogel. Upon *in situ* photo-crosslinking, the adhesive hydrogel forms a transparent bandage sealing the full-thickness wounds (yellow arrows). **B)** Image of our custommade setup with infusion setup equipped with a digital manometer for infusing saline into OcuPair sealed eyeballs in a controlled manner. **C & D)** Brust pressure measurements indicating that the OcuPair adhesive hydrogel withstands intraocular pressure (IOP) greater than normal physiological range (>50mmHg) for all the wound types. **E)** Evaluation of the adhesive strength of the OcuPair adhesive hydrogel using a modified ASTM standard, F2392-04 using porcine corneal strips as biological samples. The adhesion strength of OcuPair is compared with that of cyanoacrylate glue (Mean ± SEM, n= 8 for OcuPair and n=3 for cyanoacrylate, ** *p*<0.05, student un-paired *t*-test). **F)** Schematic of the modified test for lap shear strength measurements (ASTM F2255-05) using porcine corneal strip as biological sample and the average shear strength of OcuPair and cyanoacrylate glue (Mean ± SEM, n=3).

**Table 1:** Different corneal wounds employed and mimicking combat zone traumatic corneal injuries.

| Wound type created | Relevant traumatic condition in combat zone |
| --- | --- |
| Linear (6-8mm) | Corneal laceration caused due to blast injury or shrapnel [55] |
| 3-pronged stellate (3.5mm each arm) | Corneal puncture injury due to an irregular shrapnel or foreign body lodged in the cornea [52, 54] |
| Circular with tissue attached (4.5mm dia) | Traumatic laceration injury with lacerated tissue partially attached to the cornea. |
| Corneal Perforation (3mm dia) | Perforation or punch injury caused due to shrapnel with missing tissue [19] |
| Corneo-Scleral (6mm horizontal & 4mm vertical) | Blast injury in the corneo-scleral region [56-58] |

The first layer plugged the incision and stopped the fluid egress, and the second layer formed a hydrogel bandage acting as a barrier on the surface of the incision. The smooth hydrogel bandage was stable, without breaking when the eyes are manipulated using a cotton tipped applicator **(Please see supplementary section, Video 1 (Linear incision) and Video 2 (Stellate incision))**. Upon increasing the IOP in a controlled manner by infusing the saline into the posterior chamber of the eye globe (mimicking the increase of IOP in human setting) using our custom-made setup (**Figure 4B**), the adhesive hydrogel held and sealed the tissue for all types of incision for >60mmHg in both rabbit and porcine eyes (**Figures 4C & 4D**) which is beyond the physiological IOP range. In fact, when the saline infusion was stopped at ∼60 mmHg, the adhesive hydrogel maintained the pressure for at least 5-7 mins for linear, stellate incision and corneal perforations. We found that the adhesive hydrogel integrates well, demonstrates better sealing capabilities and withstands higher IOP in porcine corneas compared to rabbit corneal tissues for all incision types. This can be attributed to the higher corneal thickness of porcine corneas. The burst pressures for individual types of incisions are summarized in **Table S3 (supplementary information)**. At the burst point, for all full-thickness incisions, the adhesive hydrogel peeled off as a layer from the corneal surface rather than subjecting to break in the hydrogel layer, suggest that the adhesive hydrogel possesses good cohesive properties and the mechanical strength to withstand high IOP compared to commercially available ReSure® glue which demonstrated a burst pressure of ∼37 mmHg [19]. The adhesive hydrogel adhered well over the moderately dry corneal surface and could be easily peelable using common surgical forceps by pressing the corneal tissue appropriately near the hydrogel bandage edges and the tissue interface (assessed qualitatively). Additionally, we evaluated the sealing capability and burst pressure of the OcuPair hydrogel formulation made from the stability samples in *ex vivo* porcine eyes with full-thickness linear 6-8mm wound. Next, we evaluated the effect of degradation of D-MA formulation at accelerated storage conditions (40°C) on the sealing performance (burst pressure) of OcuPair adhesive hydrogel. Since, no evidence of degradation was observed in HA-MA formulation in all the storage conditions, we utilized HA-MA stored at 25°C up to 3-months and prepared the formulation (30:70, D-MA: HA-MA) with D-MA stability samples stored at 4°C, 25°C and 40°C at 3-month timepoint, to form the sealant bandage. Both D-MA stored at 4°C and 25°C for 3 months demonstrated higher burst pressures withstanding IOP of ∼90.2 ± 4.7 mmHg and ∼96.6 ± 5.0 mmHg respectively which is similar to that of regular samples (**Figure S8, Supporting information**). Whereas D-MA samples stored at 40°C, which demonstrated cleaved methacrylic acid, had lower burst pressure withstanding IOP 58.2 ± 3.7 mmHg which is lower to that of 4°C, 25°C and regular samples (p<0.001, n=4, 40°C *vs* 4°C and 25°C) (**Figure S8, Supporting information**). This further suggests that D-MA with methacrylate groups contributes to the mechanical strength to the OcuPair hydrogel and 4°C or 25°C storage conditions will be more appropriate for the developed formulation. Further, we evaluated the adhesive strength of OcuPair adhesive hydrogel with porcine corneal tissue using a modified wound closure test based on ASTM standard F2458-05 (**Figure 4E**). OcuPair adhesive hydrogel demonstrated good adhesive strength adhering and holding the two ends of the corneal tissue. When extended, it withstood the load up to ∼165.5 ± 10.2 seconds with a maximum load of ∼0.63 ± 0.11 N and reached an adhesive strength of ∼252.1 ± 50.6 kPa (n=8, mean ± SEM). On the other hand, cyanoacrylate glue (control) withstood higher load of ∼3.2 ± 0.2 N for similar time of ∼178 ± 28.2 seconds with an adhesive strength of ∼615 ± 44.8 kPa. Next, the shear strength of the OcuPair adhesive hydrogel was evaluated using a modified test base on ASTM standard F2255-05 (**Figure 4F**). The lap shear strength of OcuPair adhesive hydrogel was ∼80.5 ± 22.6 kPa, whereas cyanoacrylate had a moderately higher shear strength of ∼97.4 ± 23.1 kPa. It has been reported that cyanoacrylates demonstrate superior adhesive strength [15, 49] but it should be noted that for cyanoacrylate to adhere to the cornea, the corneal surface should be dry and we observed that after application of cyanoacrylate, it becomes hard instantly forming a rough surface making the tissue specimen hard and stiff. This effect of forming rough surface and drying of the cornea may cause aberrations in the eyelid resulting in irritation every time a patient closes the eyelids. Overall, OcuPair adhesive hydrogel demonstrates good adhesive, cohesive and mechanical properties which help it withstand IOP beyond physiological range and the pressure caused due to eye moments and frequent closure of eyelids.

### 2.7. OcuPair hydrogel adhesive secures full-thickness corneal wounds and stabilizes the cornea in a rabbit corneal injury model

We have evaluated the sealing performance of the OcuPair and its ability to stabilize various full thickness corneal and corneo-scleral wounds for a period of 5 days in a clinically relevant rabbit model. Different wound architectures were employed that collectively mimic the battlefield combat corneal injuries experienced by the wounded army personnel (**Table 1**). The 5-day endpoint was chosen, as a representative time that is required for the injured army personnel to be transported from the role 1 battalion aid station to higher role of care facilities where the ophthalmologist can access the damages and treat them appropriately [4, 6, 50]. Twenty-four hours after creation of corneal wounds (except for circular incision and corneal perforations given the severity of these wounds), the anterior chamber was re-created by gently injecting the OcuPair injectable filler gel. The injectable filler gel was able to be easily injected into the anterior chamber via a 27g cannula and can be attributed to the low viscosity and shear thinning properties hyaluronic acid in the formulation. Following the recreation of the anterior chamber, the full-thickness wounds were sealed by the adhesive hydrogel using the two-layer approach that was mentioned earlier (**Section 2.4.2 (iv)**). The adhesive hydrogel solution stays in place for at least 20 seconds after application over the cornea and can be attributed to the appropriate viscosity of the formulation. Upon photocuring with UV light for 90 seconds, a transparent hydrogel bandage was formed adhering and securing different full thickness wounds. Clinical parameters such as chamber formation, corneal epithelial edema, corneal clarity/opacity, anterior FLARE, conjunctival chemosis, Intraocular pressure (IOP) changes, signs of aqueous leakage, central corneal thickness (CCT), and macroscopic structural integrity/adherence of OcuPair hydrogel on the cornea were evaluated. The outcomes for all the different full-thickness corneal wounds at POD3 and 5 are summarized in **Table 2 and 3**.

**Table 2:** Summary of results from in vivo studies in rabbit traumatic corneal injury model for linear, stellate and corneal perforation wound groups.

| Parameters Evaluated | Full thickness corneal wound types part I |  |  |
| --- | --- | --- | --- |
|  | Linear (6-8mm) | Stellate (3 branches, 3.5mm each branch) | Corneal perforation (3mm dia) with tissue missing (evaluated only on POD3) |
| **Anterior chamber formation** | **POD3:** Score 3 (full chamber with over 80% depth) formation for all (6/6) eyes.<br>**POD5:** Score 3 for 5/6 eyes, Score 2 for 1 eye (mild shallow chamber with 50-80% depth). | **POD3:** Score 3 for 4/6 eyes and score 2 for 2/6 eyes.<br>**POD5:** Score 3 for 5/6 eyes and score 2 for 1 eye. | **POD3:** Score 3 for all (3/3) eyes. |
| **Intraocular pressure (IOP) measurements (mm/Hg)** | **POD3:** Normotensive IOP for all (6/6) eyes.<br>**POD5:** Normotensive IOP for all (6/6) eyes. | **POD3:** Normotensive IOP for all (6/6) eyes.<br>**POD5:** Normotensive IOP for all (6/6) eyes. | **POD3:** Normotensive IOP in all 3 eyes |
| **Conjunctival chemosis** | **POD3:** Score 0 (no congestion) for all (6/6) eyes.<br>**POD5:** Score 0 for 5/6 eyes, one eye had mild congestion (Score 1+). | **POD3:** Score 0 for 2 eyes, 1+ for 1 eye and 2+ for 3 eyes.<br>**POD5:** Score 0 for 3 eyes, 1+ for 2 eyes, 2+ for 1 eye. | **POD3:** Score 0 for all (3/3) eyes. |
| **Corneal Opacity/clarity** | **POD3:** Score 0 (clear cornea) for all (6/6) eyes.<br>**POD5:** Score 0 (clear cornea) for all (6/6) eyes. | **POD3:** Score 0 (clear cornea) for all (6/6) eyes.<br>**POD5:** Score 0 (clear cornea) for all (6/6) eyes. | **POD3:** Score 1+ (slight haze with iris and lens visible) for all 3 eyes |
| **Corneal epithelial edema (areas surrounding and under the hydrogel bandage)** | **POD3:** Score 0 (no edema) for all (6/6) eyes.<br>**POD5:** Score 0 for all 6 eyes. | **POD3:** Score 0 for 4 eyes, 1+ for 1 eye and 2+ for 1 eye.<br>**POD5:** Score 1+ for all (6/6) eyes. | **POD3:** Score 1+ for all (3/3) eyes. |
| **Signs of inflammation & infections (FLARE score)** | **POD3:** FLARE score 0 for all (6/6) eyes.<br>**POD5:** FLARE score 0 for 1 eye, score 1+ (mild) for 4 eyes and score 2+ (moderate) for 1 eye. | **POD3:** Score 0 for 3 eyes, 1+ for 2 eyes and 2+ for 1 eye.<br>**POD5:** Score 2+ for all (6/6) eyes | **POD3:** 1+ for all (3/3) eyes. |
| **Aqueous leakage (Siedel test)** | **POD3:** No leakage in all (6/6) eyes.<br>**POD5:** No leakage in all (6/6) eyes. | **POD3:** No leakage in all (6/6) eyes.<br>**POD5:** No leakage in 5 eyes and small leakage in 1 eye. | **POD3:** No leakage in all (3/3) eyes. |
| **Central corneal thickness (CCT) μm** | **POD3:** Corneal edema +ve in 6/6 eyes.<br>**POD5:** Corneal edema +ve in 6/6 eyes. | **POD3:** Corneal edema +ve in 6/6 eyes.<br>**POD5:** Corneal edema +ve in 6/6 eyes. | **POD3:** Corneal edema +ve in 6/6 eyes. |
| **OcuPair hydrogel bandage intactness** | The hydrogel bandage is intact and well adhered to cornea in all the (12/12) eyes both in **POD3 & 5**. | **POD3:** Hydrogel bandage intact in all (6/6) eyes.<br>**POD5:** Intact in 5 eyes and 1 eye demonstrated a crack. | **POD3:** The hydrogel bandage intact and well adhered to the cornea. |
| **Intraocular contents visibility through adhesive hydrogel** | **POD3:** Intraocular content contents are clearly visible in all 6 eyes.<br>**POD5:** Intraocular content contents are clearly visible in all 6 eyes. | **POD3:** Intraocular content contents are clearly visible in all 6 eyes.<br>**POD5:** Intraocular content contents are clearly visible in all 6 eyes. | **POD3:** Intraocular content contents are visible in all 3 eyes. |

**Table 3:** Summary of results from *in vivo* studies in rabbit traumatic corneal injury model for circular, corneo-scleral and linear (sealed with cyanoacrylate) wound groups.

| Parameters Evaluated | Full thickness corneal wound types part II |  |  |
| --- | --- | --- | --- |
|  | Circular (4.5mm Dia with tissue attached) | Corneo-Scleral (6mm horizontal and 4mm vertical) | Linear (6-8mm) treated with Cyanoacrylate glue (evaluated only on POD5) |
| **Anterior chamber formation** | **POD3:** Score 1 (shallow chamber with 30-50% depth) formation for all (6/6) eyes.<br>**POD5:** Score 2 (mild shallow chamber with 50-80% depth) for 1 eye, score 1 (shallow chamber) for 3 eyes and score 0 (no chamber) for 2 eyes. | **POD3:** Score 3 (fully formed chamber) for all 6 eyes.<br>**POD5:** Score 3 for all 6 eyes. | **POD5:** Score 1 (shallow chamber) for all 5 eyes. |
| **Intraocular pressure (IOP) measurements (mm/Hg)** | **POD3:** Hypotensive IOP for all (6/6) eyes.<br>**POD5:** Hypotensive IOP for all (6/6) eyes. | **POD3:** Normotensive IOP for all (6/6) eyes.<br>**POD5:** Normotensive IOP in 5 eyes, hypotensive in 1 eye. | **POD5:** Hypotensive IOP in all 5 eyes |
| **Conjunctival chemosis** | **POD3:** Score 0 for 2 eyes, 1+ for 1 eye and 2+ (moderate congestion) for 2 eyes.<br>**POD5:** Score 0 for 3 eyes, 1+ for 2 eyes, and 2+ for 1 eye. | **POD3:** Score 1+ for all 6 eyes.<br>**POD5:** Score 1+ for all 6 eyes. | **POD5:** Score 2+ for 1 eye and 3+ for 4 eyes. |
| **Corneal Opacity/clarity** | **POD3:** Score 0 for 4 eyes and 1+ for 2 eyes.<br>**POD5:** Score 0 for 2 eyes, 1+ for 3 eyes and 2+ for 1 eye. | **POD5:** Score 0 (clear cornea) for all 6 eyes.<br>**POD5:** Score 0 for all 6 eyes. | **POD5:** Score 1+ for 1 eye, 2+ for 3 eyes and 3+ for 1 eye. |
| **Corneal epithelial edema (areas surrounding and under the hydrogel bandage)** | **POD3:** Score 0 for 1eye, 1+ for 3 eyes and 2+ for 2 eyes.<br>**POD5:** Score 1+ for all 6 eyes. | Not suitable measure for this type of incision | **POD5:** Score 1+ for 2 eyes and 2+ for 3 eyes. |
| **Signs of inflammation & infections (FLARE score)** | **POD3:** FLARE score 0 for 1 eye, 1+ for 1 eye and 2+ for 4 eyes.<br>**POD5:** FLARE score 1+ for all 6 eyes. | **POD3:** Score 0 for all 6 eyes.<br>**POD5:** Score 0 for all 5 eyes, 1+ for 1 eye. | **POD5:** Score 3+ (marked FLARE with iris and lens details are hazy and hyphema) for all 5 eyes. |
| **Aqueous leakage (Siedel test)** | **POD3:** Aqueous leakage in 1 eye.<br>**POD5:** Aqueous leakage in 2 eyes. | **POD3:** No leakage in all 6 eyes.<br>**POD5:** No leakage in 5 eyes and leakage in 1 eye. | **POD5:** Aqueous leakage in all 5 eyes. |
| **Central corneal thickness (CCT) μm** | **POD3:** Corneal edema +ve in 6/6 eyes.<br>**POD5:** Corneal edema +ve in 6/6 eyes. | Not suitable measure for this type of incision | **POD5:** Not measured due to opaque cyanoacrylate layer on cornea. |
| **OcuPair hydrogel bandage intactness** | **POD3:** Intact in 5 eyes, partial peeling and dislodgement in 1 eye.<br>**POD5:** Intact in 4 eyes, partial dislodgement in 2 eyes. | **POD3:** Hydrogel intact in all 6 eyes.<br>**POD5:** Intact in 5 eyes and 1 eye demonstrated breakage and partial dislodgement. | **POD3:** The cyanoacrylate glue was completely dislodged in 3 eyes and partially attached in 2 eyes. |
| **Intraocular contents visibility through adhesive hydrogel** | **POD3:** Clearly visible in all 4 eyes, visible with slight haze in 2 eyes.<br>**POD5:** Visible with slight haze in all 6 eyes. | **POD3:** Intraocular content contents are clearly visible in all 6 eyes.<br>**POD5:** Intraocular content contents are clearly visible in all 6 eyes. | **POD3:** Contents are not visible through cyanoacrylate layer. |

We did not observe any signs of swelling or significant macroscopic structural integrity changes of the hydrogel bandage (**Figure 5**). Most of the eyes (>95%) in linear, stellate corneal perforation, corneo-scleral wound groups demonstrated intact hydrogel bandage (**Figure 5A and 5B**). Further, the intactness of the hydrogel bandage was evaluated by gently manipulating the cornea using sterile cotton swap. Two eyes in linear incision group at POD5 demonstrated minor peeling at the edges of the hydrogel bandage in the superior cornea and 1 eye each at POD5 in both stellate incision and corneo-scleral wound groups demonstrated a crack (**Figure 5B (22)**). This can be attributed to frequent eyelid closure and eye moments that can exert stress over the hydrogel layer. In the circular incision group, we observed partial peeling dislodgement with more than 50% adherence to the cornea (estimated qualitatively using visual inspection under the surgical microscope) in 3 eyes (1 at POD3 and 2 at POD5). Given the architecture and severity of the wound which may exert more pressure combined with frequent eyelid moments, such partial dislodgement at superior quadrant of the cornea can be expected (**Figure 5B (11)**). The slit-lamp microscopy examination demonstrates that the adhesive hydrogel maintained its transparency with intraocular contents clearly visible through OcuPair adhesive hydrogel bandage in more than 95% of the eyes in all the wound groups except for the circular wound. Here, the intraocular contents are visible with some degree of haziness when viewed through the central corneal button but clearly visible through the hydrogel layer and the corneal tissue beneath it in the areas other than the central corneal buttons (**Figure 5B (4 & 10)**).

**Figure 5:**
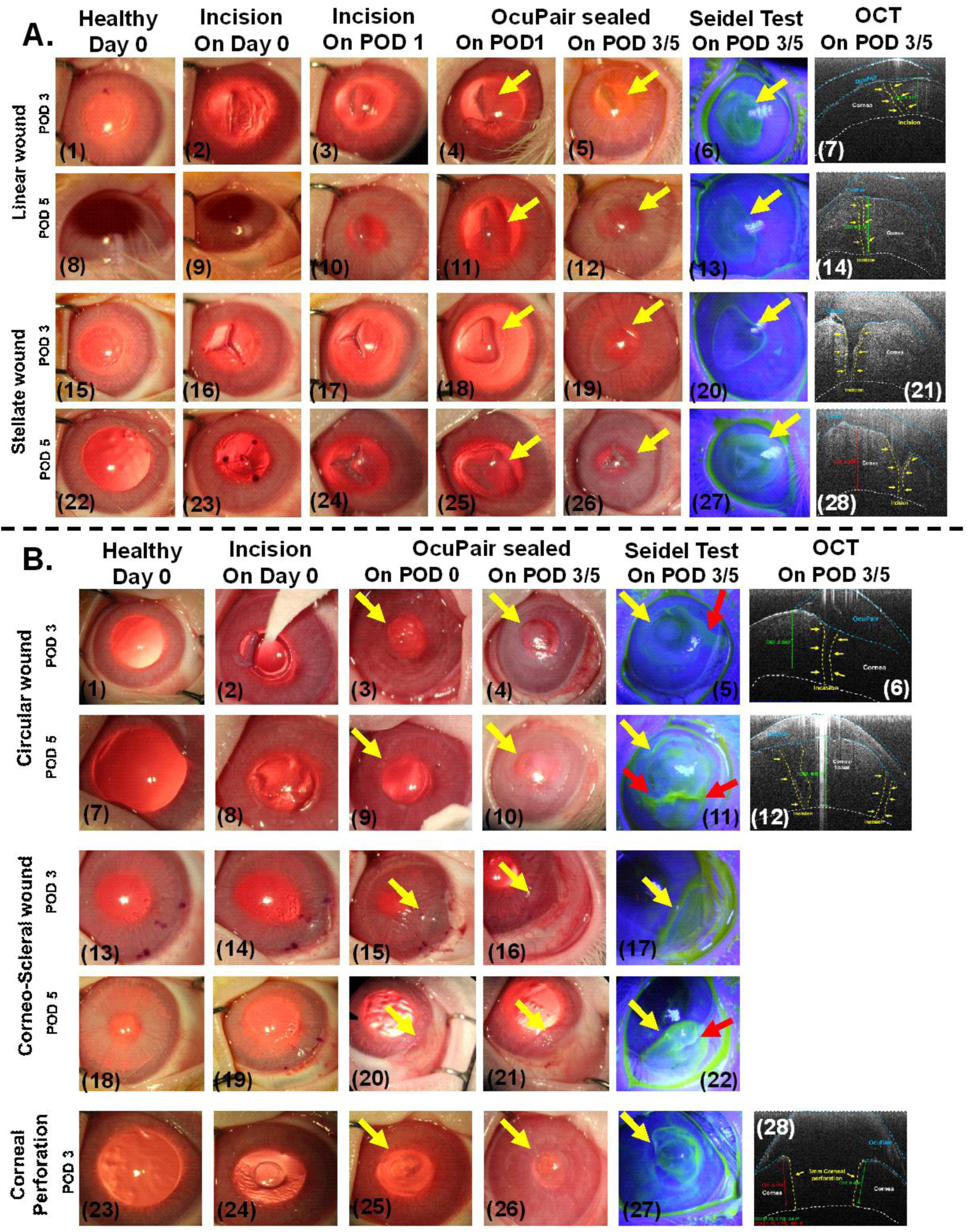
*In vivo* evaluation of OcuPair in a rabbit corneal injury model. **A)** Representative photographic, slit lamp and optical coherence tomography (OCT) images of rabbit eyes with full-thickness linear and stellate wounds sealed with OcuPair adhesive hydrogel at POD3 and POD5 timepoints. **B)** Representative photographs and OCT images of rabbit eyes with circular, corneo-scleral wounds and corneal perforation sealed with OcuPair. The OcuPair adhesive hydrogel seals the full-thickness wounds forming a transparent hydrogel bandage adhering to the cornea and stabilizing the eye until POD5. Yellow arrows indicate the transparent hydrogel bandage, and red arrows indicate the disruption of glue integrity and leakage from the incision site.

More than 90% of the eyes with wounds sealed with OcuPair adhesive hydrogel in linear, stellate, corneal perforations and corneo-scleral groups demonstrated fully formed chamber (over 80% depth) with a score of 3 in POD3 and POD5 timepoints (**Figure 6B**). A mild shallow chamber (50-80% depth) with a score of 2 was observed only in ∼8% of the eyes especially in POD5 timepoint (**Figure 6B**). Further, Seidel test revealed that, all the eyes with fully formed chamber and few eyes with moderately shallow chamber demonstrated no leakage from the incision site. Only the eyes that had a crack (1 eye in stellate group and 1 eye in corneo-scleral group) demonstrated leakage from the incision site at POD5 timepoint (**Figure 5B, (22)**). All the eyes in these groups (∼98%) demonstrated normotensive IOP whereas only 1 eye in corneo-scleral group at POD5 demonstrated hypotensive IOP compared to baseline IOP which also demonstrated leakage. In the circular incision group, ∼80% of the eyes demonstrated shallow chamber (30-50% depth) with a score of 1(**Figure 5B, (4 & 10)**). Approximately 12% of the eyes demonstrated mild shallow chamber (50-80% depth) with a score of 2 and ∼8% (2 eyes at POD5) of the eyes demonstrated no chamber formation (**Figure 6B**). The Siedel test demonstrated that 1 eye at POD3 and 2 eyes (that have no chamber) at POD 5 demonstrated aqueous leakage (**Figure 5B (5 & 11)**). These results are consistent with the eyes where partial dislodgement of OcuPair adhesive hydrogel bandage was observed. Furthermore, the severity of circular incision should also be considered as a greater number of eyes that demonstrated hydrogel bandage dislodgement and leakage with shallow chamber were observed at POD5 timepoint.

**Figure 6:**
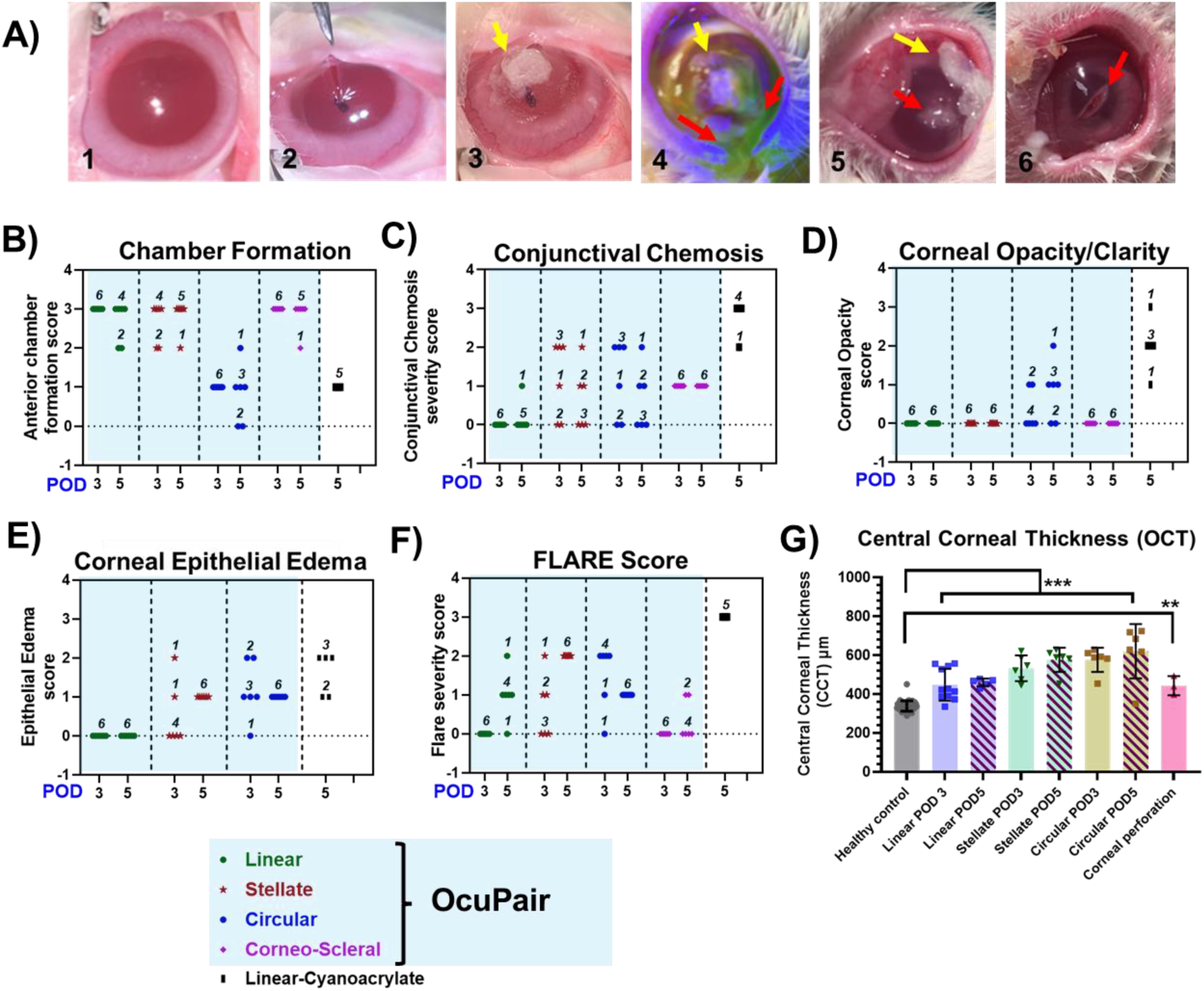
A) Representative images of rabbit cornea with full thickness corneal wound sealed with cyanoacrylate glue. Image #3 wound sealed with cyanoacrylate glue on POD1, images 4-6 were taken on POD5, demonstrating leakage form the wound and dislodgement of the cyanoacrylate glue from the corneal surface. Note in image 5, plugging of the incision with inflammatory secretions indicated by red arrow and after removing the secretions and cleaning the cornea, revels reddening of the tissue at the incision site indicated with red arrow (image 6). **B-G) Assessment of performance and efficacy of OcuPair via clinical benchmarks from slit-lamp examination at POD 3 and POD 5: B)** Anterior chamber formation, **C)** Conjunctival chemosis score, **D)** Corneal opacity/clarity score, **E)** Corneal epithelial edema, **F)** FLARE score and **G)** Central corneal thickness from OCT examination (data represented as mean ± SEM, *** p <0.001 & ** *p* <0.05, One-way ANOVA). **Note:** The numbers indicated over the data points in graphs D-F represent the number of eyes with a particular score.

The eyes in linear and corneal perforation wound groups demonstrated no conjunctival chemosis with a score of 0. Only 1 eye in the linear wound group had mild conjunctival chemosis at POD5 (**Figure 6C**). In the stellate and corneo-scleral wound groups, mild conjunctival chemosis with a score of 1+ was observed in most of the eyes. Only a few had moderate conjunctival chemosis with a score of 2+ were observed (**Figure 6C**). In the circular incision group, many eyes demonstrated moderate conjunctival chemosis with a score of 2+ (**Figure 6C**). All the eyes in linear, stellate and corneo-scleral wound groups both in POD3 & POD5 demonstrated good corneal clarity in the tissue beneath adhesive hydrogel bandage and area surrounding it with a corneal opacity score of 0 suggesting that OcuPair does not induce any significant corneal toxicity that leads to opacification of the cornea (**Figure 6D**). In the corneal perforation group, all eyes had very mild opacity with a score of 1+. In the circular incision group ∼50% of the eyes demonstrated clear cornea. Mild to moderate (score 1+ to 2+) corneal opacity was observed in another 50% of the eyes mostly at POD5 timepoint (**Figure 6D**). This loss of transparency was only observed in the central corneal buttons area and can be attributed to the severity of the wound and the partial attachment of the tissue may not be enough for supply of nutrients necessary for tissue viability (**Figure 5B, (4 & 10)**). Corneal epithelial edema in the corneal tissue under the hydrogel bandage and its surrounding areas was not observed in the linear wound group (**Figure 6E**). The corneal perforation group demonstrated mild corneal edema with a score of 1+ and was observed in all 3 eyes. Few eyes in stellate and circular wound groups also demonstrated no epithelial edema but many eyes (∼80%) demonstrated mild to moderate corneal epithelial edema (**Figure 6E**) and can be attributed to severity of the created wounds.

Any signs of anterior chamber inflammation and infections were evaluated using flare score (**Figure 6F**). Most of the eyes (∼87%) in all the wound groups at POD3 demonstrated no flare with a score of 0 (**Figure 6F**). At POD5, most of the eyes in all the wound groups had faint flare (score of 1+). Moderate flare (score of 2+) with iris and lens details clear was more pronounced in circular incision group (**Figure 6F**). OCT imaging studies demonstrated that the OcuPair adhesive hydrogel is adhered to the corneal surface both in POD3 and POD5 eyes sealing and stabilizing the full-thickness wounds and maintaining the corneal architecture and with good anterior chamber in most of the eyes in all the wound groups except for the ones that have shallow or no chamber (**Figures 5A (7, 14, 21 & 28) & 5B (6, 12, 28)**). Further corneal thickness measurements demonstrate evidence of stromal edema resulting in significant CCT thickness in all the wound groups compared to their respective contralateral healthy controls. The increase in CCT due to corneal edema can be attributed to the creation of the wound that can include trauma to the cornea (**Figure 6G**).

As a control for OcuPair adhesive hydrogel, we evaluated the performance of Histoacryl, a type of cyanoacrylate glue that can be used to treat corneal perforations and descemetoceles [13]. A linear full thickness laceration (5-6mm) was employed (**Figure 6A, 2**) and the wound was sealed using cyanoacrylate glue. Upon application of the cyanoacrylate glue on the incision, it solidified instantly forming a hard and opaque layer with irregular/rough surface on the cornea (**Figure 6A, 3 yellow arrow**). To avoid eye lid aberration and discomfort we used contact lens, but due to irregular protrusions and rapid eye blinking in rabbits, the lens was dislodged within 24 hrs. The glue was opaque, and the intraocular contents were not visible in slit lamp evaluation (**Figure 6A, 3-6**). At POD3, we observed dislodgement/peeling (∼40-50%, qualitatively) of the glue from the corneal surface at the edges and by POD5 all eyes had dislodged glue with open incision plugged with white/yellowish inflammatory secretions (**Figure 6A, 5 red arrows**). We did not observe such inflammatory secretions in any of the OcuPair groups suggestive of the effect of cyanoacrylate glue. Slit lamp examination revealed mild to moderate corneal toxicity and edema with a score of 2+ to 3+ for all the eyes (5/5) at the near vicinity/edges of the glue suggests unfavorable reaction of the corneal tissue to the cyanoacrylate. Evidence of corneal reddening (mild, at the incision site, possibly due to the inflammatory secretions plugging the incision) corneal epithelial edema and stromal edema was more pronounced in this group compared to our OcuPair groups (**Figure 6E**). Due to the formation of the rough surface, aberrations in corneal eye lids were observed. The Seidel test demonstrated aqueous leakage from the incision site in all the eyes (**Figure 6A, 4 red arrows**). A flare score of 3+ was given to all the eyes in this group due to the above reasons (**Figure 6F**). Even though the peripheral cornea appeared to be healthy, corneal reddening (mild) was a concern (**Figure 6A, 6 red arrows**).

### 2.8. OcuPair demonstrates favorable histological outcomes and biocompatibility in corneal tissue compared to cyanoacrylate glue controls

The sections were stained for hematoxylin and eosin to evaluate the corneal toxicity and immune response of OcuPair adhesive hydrogel. The healthy tissue revealed normal corneal architecture, multi-layered corneal epithelium and stroma with normal keratocytes (**Figure 7A, 1 & 6**). The cornea sealed with cyanoacrylate control revealed inflammation at the incision site plugging the incision at POD5 timepoint (**Figure 7A, 2**). The cyanoacrylate adhesive was missing from corneal surface in all the eyes suggesting their dislodgement. These findings were consistent with our clinical assessment where inflammatory secretions plugging the corneal wound (**Figure 7A, 2**). Thinning of corneal epithelial layer at the area where the cyanoacrylate was applied on the cornea suggests unfavorable reaction of corneal epithelium to the cyanoacrylate (**Figure 7A, 2**). The stromal tissue in the cyanoacrylate demonstrates larger vacuoles like structures suggests stromal edema resulting partly from the trauma due to incision and partly due to the unfavorable reaction to cyanoacrylate (**Figure 7A, 2 & 7, yellow arrows**). Additionally, this control group also demonstrated a higher number of inflammatory cells (rounded) in the stromal tissue and the near vicinity of the wound site suggests inflammation (**Figure 7A, 7**). In the OcuPair sealed group, the adhesive hydrogel was intact and adhered to the corneal surface in all the wound groups consistent with our clinical observations. Increase in total corneal thickness was noted (qualitatively) similar to that of cyanoacrylate group suggests the effect of trauma due to the creation of the wound. Interestingly, stromal edema was less pronounced in the OcuPair group, particularly near the areas that are in close contact with hydrogel compared to that of cyanoacrylate group suggesting that OcuPair may not induce stromal edema (**Figure 7, 3 – 5 & 8 – 10**). All the OcuPair treated eyes have some degree of keratocyte activation or presence of inflammatory cells but less severe when compared to eyes treated with cyanoacrylate control (**Figure 7, 8 – 10**).

**Figure 7:**
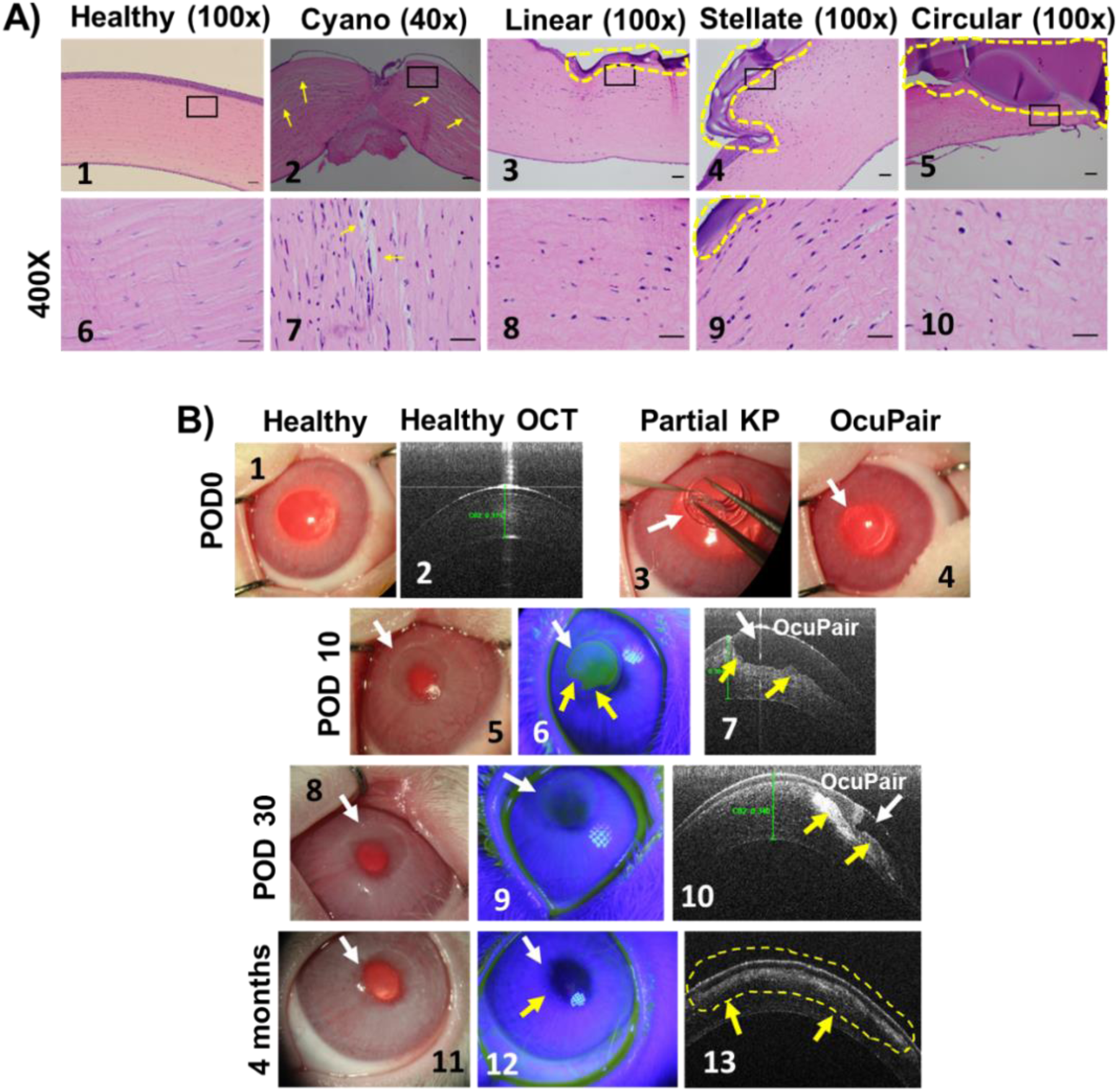
Toxicity and long-term biocompatibility assessment of OcuPair adhesive hydrogel. **A)** Representative hematoxylin and eosin histopathology images at POD5 timepoint from **(1 & 6)** Healthy cornea, **(2 & 7)** Cyanoacrylate (Cyano) demonstrating significant stromal edema (yellow arrows) and inflammatory cells. **(3 & 8)** Linear wound group with OcuPair adhesive hydrogel intact on the corneal surface (yellow dotted lines). **(4 & 9)** Stellate group with OcuPair (yellow dotted lines) and **(5 & 10)** Circular wound group sealed with OcuPair (yellow dotted lines). Scale bar: 100μm. **B)** Assessment of long-term biocompatibility of OcuPair in corneal tissue. Representative images of healthy cornea and OCT **(1 & 2)** before partial keratoplasty (partial KP) defect in the cornea. **(3 & 4)** partial KP defect was created and treated with OcuPair adhesive hydrogel (white arrows). **(6 – 7)** At POD 10 the native cornea surrounding the OcuPair hydrogel is clear and did not show any signs of corneal toxicity at the interface. The OCT image **(7)** demonstrates that hydrogel (white arrow) is week adhered to stroma (yellow arrows). **(8 – 10)** Images at POD 30, demonstrating signs of epithelization over the OcuPair layer and OCT analysis demonstrates stromal integration (yellow arrows). **(11 – 14)** 4 months after OcuPair treatment demonstrating significant stromal integration into the OcuPair layer (Yellow arrows and dotted lines).

The GLP toxicity studies were conducted in healthy rabbits and the results suggest that both injectable filler gel and the individual components of the adhesive hydrogel (D-MA, HA-MA and Irgacure2959) in pre-hydrogel solution form, was well tolerated in all the animals and did not exhibit any noticeable macroscopic toxicity or any signs of corneal and anterior chamber infections or inflammation during clinical examination. Slit lamp examination revealed adhesive hydrogel deposit at the inferior anterior chamber with no evidence of anterior chamber uveitis in all the animals. One animal has anterior lens capsule opacity at the inferior region on day 7 but resolved completely before day 29. The approach of subconjunctival route was adopted as these tests were conducted in healthy rabbits where there is no provision to anchor the adhesive hydrogel to the corneal surface and subconjunctival tissue is in close contact with peripheral cornea [51].Subconjunctival injection of adhesive hydrogel solution followed by photo-crosslinking resulted in hydrogel deposit which persisted till day 30 in the subconjunctival space and was well tolerated. Minimal to mild conjunctivitis was observed in all the animals in this group between days 3 and 7 but eventually resolved by day 10 and may be attributed to the space-occupying nature of the depot injection within the subconjunctival space. Histological analysis identified the subconjunctival adhesive hydrogel in most animals demonstrated tissue reaction consisted of a minimal to mild fibrous capsule with macrophages and multinucleated giant cells at the interface between the capsule and gel, which was considered normal for a degradable material at a 30-day termination interval. In conclusion, administration of OcuPair or its components by bilateral ocular injections into different ocular compartments or tissues of NZW rabbits was well-tolerated with no signs of ocular toxicity after 30 days post-injection.

To evaluate the long-term corneal toxicity and biocompatibility of OcuPair adhesive hydrogel, a partial keratoplasty (PKP) was made to create a defect in the corneal tissue to hold the adhesive hydrogel in place in the tissue for longer period. This serves as a mode to evaluate the effect of hydrogel on corneal stroma and epithelium. After filling the defect with adhesive hydrogel solution followed by photo-crosslinking with UV light for 90 seconds resulted in a transparent hydrogel firmly adhered to the corneal defect (**Figure 7B, 4**). Ten days after implantation, the adhesive hydrogel was transparent with smooth surface, further slit-lamp examination revealed that the surrounding tissue was transparent and non-inflamed (**Figure 7B, 5**). OCT examination also demonstrates that OcuPair hydrogel is intact, well adhered to the corneal stroma and maintains the corneal architecture with signs of corneal epithelial growth. (**Figure 7B, 7**). At POD30, the adhesive hydrogel is still transparent with no signs of inflammation or infections in the corneal tissue surrounding the hydrogel. Further, no evidence of corneal opacity of the native cornea at the hydrogel edges further confirms its biocompatibility. The Siedel test demonstrates some degree of corneal epithelial growth over the hydrogel surface. OCT imaging revealed that signs of hydrogel degradation and stromal integration or regeneration at the interface of hydrogel and corneal tissue. Four months after implantation, the corneal tissue did not show any signs of inflammation or opacification, and OCT imaging demonstrates degradation of OcuPair adhesive hydrogel, stromal integration and epithelium growth with minimal hyper reflectivity suggestive of hydrogel’s biocompatibility.

## 3. Conclusions

Unlike surgical corneal wounds, battlefield related, and traumatic corneal injuries are often complex, challenging and are highly susceptible to infections, inflammation and reduced intraocular pressure [52]. If not treated promptly, proper ocular physiology and nutrient supply is greatly impacted resulting in poor visual outcomes or necrosis, necessitating enucleation of the eye [53, 54]. Hence the need for novel solutions that can stabilize the wounds quickly, preserving ocular components till more permanent solutions are planned by ocular traumatic surgeons at the operation theater. Aiming the latter, we have developed and validated an OcuPair hydrogel system that addresses both hypotony and seals full thickness wounds at the combat zone role of care without the requirement of sophisticated instrumentation.

The components of OcuPair have been chosen so that they are compatible with ocular tissues, that include a non-toxic hydroxyl PAMAM dendrimer and HA that is already a component of cornea. The adhesive hydrogel components (D-MA and HA-MA) developed here exhibits a high degree of methacrylate substitution without affecting its original properties such as solubility, shear-thinning and mucoadhesiveness but contributing to gels with high crosslinking densities. The adhesive hydrogel formulation developed in this work has: (i) appropriate viscosity that upon application that enables it to stay in place over the corneal tissue allowing time for medical personnel to photocure the applied layer. (ii) photo-crosslinks within 90 seconds of low powered UV light to form transparent and smooth hydrogel bandages securing full thickness corneal and corneo-scleral wounds the mimics combat zone ocular injuries. (iii) is flexible and possesses good mechanical and adhesive properties withstanding high intraocular pressure (>60mmHg) beyond normal physiological IOP *ex-vivo*.

In a rabbit model of corneal injury, the OcuPair adhesive hydrogel sealed and stabilized complicated full thickness wounds of various architectures forming a transparent hydrogel bandage with smooth surface upon photo-crosslinking securing and maintaining the corneal architecture. The hydrogel bandages adhered well to the corneal surface creating a barrier protecting the open wounds without any signs of leakage in majority of animals (∼98%) until 5 days after application and allowing visualization of intraocular contents through them. This will allow the ocular surgeons to evaluate the intraocular damage. The adhesive hydrogel bandage demonstrated favorable clinical benchmarks maintaining the corneal clarity and IOP’s in normotensive range. OcuPair hydrogel did not induce any significant signs of corneal toxicity, conjunctival chemosis, corneal epithelial edema, and with very minimal flare compared to eyes sealed with cyanoacrylate glue. The OcuPair hydrogel bandage did not induce significant discomfort while eyelid closure in eyes sealed with OcuPair, as opposed to cyanoacrylate treated eyes with evidence of aberrations and reddening of inner surface of the eyelids.

Additionally, histological assessment demonstrated that both components of OcuPair (injectable filler gel and adhesive hydrogel) did not induce any significant signs of corneal toxicity with minimal inflammatory cells accumulation in the wound site. GLP toxicity studies also demonstrated all the individual components of OcuPair did not induce sub-clinical intraocular toxicity or any significant adverse effects even at 30 days post implantation of adhesive hydrogel into subconjunctival space in healthy rabbits. Further, long term toxicity studies in healthy rabbits with partial keratoplasty demonstrates that OcuPair adhesive hydrogel and its structural components were biocompatible with the corneal stroma and epithelium and did not illicit any significant signs of corneal toxicity.

Future IDE enabling studies evaluating this platform for various ocular applications will lead towards clinical translation for improved platforms at low-cost settings.

## 4. Experimental Section

### 4.1. Chemical and reagents

Hydroxyl-functionalized ethylenediamine core generation four PAMAM dendrimers (G4-OH; clinical grade; 64 end-groups) was purchased from Dendritech Inc. Sodium hyaluronate with average molecular weights of ∼700kDa (for injectable filler gel) and ∼42kDa (for adhesive hydrogel) were purchased from Lifecore Biomedical (Chaska, MN, USA). Methacrylic acid, Methacrylic anhydride, and 4-Dimethylaminopyridine (DMAP), 1-(3-Dimethylaminopropyl)-3-ethylcarbodiimide hydrochloride (EDC.HCL), Triethylamine (TEA), Dowex 50W X8 Ion Exchange Resin, Sodium chloride (NaCl), anhydrous dimethylacetamide (DMAc), anhydrous dimethyl sulfoxide (DMSO), were purchased from Sigma-Aldrich (St. Louis, MO, USA). ACS grade DMAc, methanol, ethanol, HPLC grade water, acetonitrile and methanol were purchased from Fisher Scientific. Dialysis membrane (MW cut-off 1000 & 2000 Da) was obtained from Spectrum Laboratories Inc. (Rancho Dominguez, CA, USA). Pellicon 3 (3kDa and 5kDa) membrane cartridge for tangential flow filtration (TFF) were purchased from EMD Millipore, USA.

Proton NMR spectra of the final conjugates as well as intermediates were recorded on a Bruker (500 MHz) spectrometer using commercially available DMSO-*d_6_* solvent. Proton chemical shifts were reported in ppm (δ) and tetramethylsilane (TMS) used as internal standard. All data were processed using ACD/NMR processor software (Academic Edition).

### 4.2 Synthesis of OcuPair adhesive components

#### 4.2.1 Synthesis of Dendrimer-Methacrylate (D-MA) conjugates

For the synthesis of D-methacrylate (6), we received the Generation 4, hydroxyl-terminated Pharma-grade PAMAM dendrimer from Dendritech (Midland, MI), which has a generational purity of over 95% as confirmed by HPLC. The dendrimer methanolic solution was first evaporated thoroughly in a pre-weighed round-bottom flask, and the weight was recorded. The G4-OH (10 g, 0.70 mmol) dendrimer was dissolved in anhydrous *N*, *N*-dimethylacetamide. The reaction mixture was sonicated to form a clear solution, followed by the addition of 30 mole-equivalents of DMAP (2.56 g, 21 mmol). The reaction mixture was then stirred at room temperature for 20 minutes. Methacrylic anhydride (2.69 g, 17.5 mmol, 25 mole equivalents) was added dropwise to the reaction mixture. To prevent polymerization in the reaction, 1 mole equivalent of hydroquinone monomethyl ether (MeHQ) (0.084 g, 0.7 mmol) was added, and the reaction mixture was stirred in the dark for 48 hours. the D-MA reaction mixture was purified using tangential flow filtration (TFF) equipped with Pellicon 3 (5kDa) membrane cartridge (EMD Millipore, USA). The TFF was performed for 12 hours with DMA followed by 12 hours with water to remove DMA solvent. The resultant water layer was lyophilized to obtain fluffy white powder for D-MA (6.1g). ^1^H NMR (DMSO-*d_6_*) δ 1.9 (d, *-CH_3_* of methacrylate), 2.11-3.49 (m, –C*H_2_* protons of G4-OH), 4.02 (t, –C*H_2_*OC=O protons, G4-OH), 4.73 (bs, –O*H* protons of G4-OH), 5.6 & 6.0 (bs –CH*_2_* protons of methacrylate), 7.82-8.07 (m, amide protons of G4-OH). The above reaction procedure was repeated thrice to obtain 5 different batches of D-MA with consistent methacrylate substitution.

#### 4.2.2 Synthesis of Hyaluronic Acid-Methacrylate (HA-MA) conjugates

Prior to the synthesis of HA-MA, the sodium salt of hyaluronic acid (HA) was converted to tetrabutylammoninum salt (HA-TBA) in the first step, so that it could be solubilized in DMA/DMSO (3:1) for efficient substitution reaction. To make HA-TBA, sodium hyaluronate (NA-HA) was dissolved in ultra-pure DI water at 2% (w/w) and the ion exchange was done by adding Dowex 50WX8 proton exchange resin (3 g resin per 1g HA) with vigorous stirring for 5 h. The resin was filtered off using Whatman filter paper and the filtrate was titrated to a pH of 7.1-7.5 with tetrabutylammoninum-hydroxide (TBA-OH). The product was lyophilized at –80°C to obtain an off-white floppy solid material that was then dissolved in DI-H*_2_*O and subjected to water dialysis (membrane MWCO = 2 kDa) to remove excess TBA-OH. The resultant water layer was then subjected to lyophilization and stored at –20°C until used. The ion exchange from was confirmed using ^1^H NMR analysis in D*_2_*O. ^1^H NMR (D*_2_*O-*d_2_*) 0.91 (m, –*CH_2_* protons of TBA), 1.35 (m, –*CH_2_* protons of TBA), 1.4 (t, –*CH_2_* protons of TBA) and 3.23 (t, –*CH_2_* protons of TBA).

In the second step, HA-TBA (6g, 0.084 mmoles) was first dissolved in 80mL DMSO and the solution was sonicated under nitrogen environment for 15-20 minutes. *N-N* dimethylacetamide (DMA) 320 mL was added to the reaction mixture and the reaction mixture was stirred for 20-30 minutes to get HA-TBA dissolved completely. To the reaction mixture, triethylamine (3.5mL, 25.3 mmoles) was added. The reaction mixture was stirring for another 30 minutes, followed by the addition of methacrylic anhydride (2.1mL, 14.3 mmoles) and was stirred in dark for 48 hours. After 48 hours, the reaction mixture was purified using TFF employed with 5kDa membrane cartridge to remove unreacted methacrylic anhydride and side products followed by TFF purification in water to remove DMA solvent. The resultant water layer was lyophilized to obtain TBA-HA-MA solid, and the methacrylate substitution was confirmed using ^1^H NMR. In the third step, the solid was re-dissolved in NaCl solution (0.5 gm NaCl per 100 mL of H_2_O) and stirred for 1 hour to facilitate ion exchange from TBA to NA. The resultant solution was dialyzed against ultrapure DI water to remove excess NA and TBA salts using TFF. The final eater layer was lyophilized to obtain white fluffy powder of HA-MA (6.8g). The removal of TBA and conjugation of methacrylate group was determined using ^1^H NMR. ^1^H NMR (D_2_O-*d*_2_) δ 1.85 (m, *-CH*_2_ protons of methacrylate), 1.98 (s, *-OCH*_3_ protons of HA), 3.0-4.7 (mm, *-CH*_2_ and *-CH* protos of HA), 5.6 and 6.0 (bs, *-CH*_2_ protons of methacrylate). The above reaction procedure was repeated thrice to obtain 3 different batches of HA-MA with consistent methacrylate substitution.

### 4.3. Formulation and characterization of injectable filler hydrogel

#### 4.3.1. Formulation of injectable filler gel

The injectable filler gel was formulated using low molecular weight (∼700 kDa) sodium hyaluronate (Na-HA) (Lot #025347, Lifecore Biomedical, MN, USA) in phosphate buffer solution at different concentrations of 1.5%, 2% and 3%. Na-HA (1.5g, 2g and 3g) were dissolved in 100 mL phosphate buffer and was stirred at room temperature (∼25°C) for 12 hrs at 300 rpm. The resultant viscous solutions were transferred into 50mL centrifuge tubes and centrifuged at 4°C for 10 mins at 2000 rpm to remove bubbles. The obtained clear solution was sterilized using autoclave (20 mins, 120°C and 15 psi pressure). The injectable filler gel was stored at 4°C and used periodically by sampling under sterile conditions. The injectable hydrogel was loaded into a 3 mL syringes (BD) and fitted with a disposable, sterile 27G or 30G anterior chamber cannula (Accutome, Malvern, PA) and maintained sterile for *in vivo* experiments.

#### 4.3.2. Characterization of injectable filler gel

##### (a) Viscosity measurements

The viscosity of the injectable filler gel (Component 1) was measured using two methods ***(i) Dynamic rheology method:*** dynamic or complex viscosity was measured using a horizontal rheometer (AR20000, TA instruments, USA) fitted with parallel plates (25mm diameter, serrated), with a 0.8 mm gap with controlled hydrated atmosphere at 37°C to mimic physiological conditions. A frequency sweep (100 – 0.1 Hz) with constant strain of 5% was maintained and the complex/dynamic viscosity was noted at 0.1Hz by loading ∼300 µL of the sample between the plates. ***(ii) Capillary viscometer method:*** Apparent viscosity at constant shear rate was measured the apparent using a capillary micro-viscometer (microVISC, RheoSense, USA). The conditions for the viscosity measurements were temperature = 25°C, sampling volume =100µL, prime volume = 10µL and the shear rate = 100 Hz.

##### (b) Osmolality and pH measurements

For the osmolality analysis, 25 µL of each sample at each time point was taken and diluted to 50 µL with water to reduce the viscosity. The concentration of the solution was around 95mg/mL. The vapor pressure osmometer (Wescor) was calibrated according to the guidelines provided in the user manual. A 10 µL of the solution was injected into the osmometer and readings were recorded. For the pH analysis of the samples, the calibration of pH meter was done by running the pH buffers at pH 4.0, 7.0, and 10.0. Once the pH meter is calibrated, the readings are recorded.

##### (c) Endotoxin content

Endotoxin content of the injectable filler gel was evaluated using ToxinSensor™ chromogenic LAL endotoxin assay kit (L00350, GeneScript, NJ, USA). Total endotoxin content was calculated using the calibration graph according to the manufacturer’s instructions.

### 4.4 Formulation, characterization, and stability analyses of adhesive hydrogel and its components (D-MA and HA-MA)

#### 4.4.1. Formulation of adhesive hydrogel solution

The adhesive hydrogel was formulated by using dendrimer component (D-MA) solution and hyaluronic acid component (HA-MA) at a ratio of 7:3 with catalytic amount of photo-initiator Irgacure 2959. The individual component solutions, D-MA (300mg/mL) and HA-MA (190mg/mL) were formulated in 10mM phosphate buffer and Irgacure 2959 (1g/mL) was formulated in DMSO. The individual solutions were centrifuged at 2000 rpm for 1 min at 4°C to remove air bubbles and stored at 4°C under dark until use. Before mixing the individual solutions were loaded into respective 1 mL glass syringes (Gerresheimer, Düsseldorf, Germany) with Leur-lock tips (D-MA 300μL in D-MA syringe, HA-MA 700μL + 6μL Irgacure 2959 in HA-MA syringe). The two solutions were mixed by connecting the two syringes using Luer-lok connector and pushing the pistons back and forth to enable the contents of the two syringes to mix evenly and finally the entire mixed solution was pushed to one single syringe and fitted with 27G anterior chamber cannula. A OcuPair kit was designed and assembled that consists of (i) A prototype final delivery device for injectable hydrogel, (ii) A prototype final delivery devices for adhesive hydrogel components and mixing and (iii) A detailed directions of use considering the end user feedback. A prototype of the final delivery device was designed to meet optimal use requirements at the end users (**Figure S9, Supporting information**)

#### 4.4.2. Characterization of adhesive hydrogel

##### (i) Viscosity measurements

The viscosity of the individual solutions (D-MA and HA-MA) and the mixed solution were assessed using capillary micro-viscometer (microVISC, RheoSense, USA). The conditions for the viscosity measurements were temperature = 25°C, sampling volume =100µL, prime volume = 10µL and the shear rate = 100 s^-1^.

##### (ii) Qualitative assessment of gelling, flexibility, and transparency of the adhesive hydrogel

Qualitative assessment of gelling and formation of adhesive hydrogel was performed by instilling ∼150μL of mixed pre-polymer adhesive hydrogel solution into 12X 5 mm TEM rubber molds followed by cobalt blue UV light pen 365nm (Jaxman Electronics), distance 5-7 cms from the light to the solution surface. The gelling time and confirmation of gel formation was assessed using a timer and manually manipulating the formed gel using spatula. The flexibility of the formed adhesive hydrogels was by formation of the liquid to the gel using a timer. The flexibility was qualitatively assessed by holding the formed gel strips with two forceps and manipulating the gels by twisting and folding. The transparency of the gel was evaluated by adding 100 µL of the pre-polymer solution into a 96 well plate and cured for 90S and the OD was read at 480 nm using plate reader and compared it with 100µL of water.

##### (iii) Rheological analysis of adhesive hydrogels

Rheological experiments were carried out on a horizontal rheometer (AR20000, TA instruments, USA) using the parallel plates with controlled hydrated atmosphere at 37°C in the oscillatory shear mode as previously reported [37]. Briefly, the real time gelling and gel formation were evaluated using dynamic time sweep with controlled strain of 1% using a parallel plate (15 mm diameter, serrated), 1.0 mm gap between the plates. Approximately 100 µL of adhesive hydrogel solution was loaded between the plates and the gelling time or the crosslinking time was evaluated by illuminating the UV light on to the pre-polymer solution. The storage modulus G’, loss modulus G”, dynamic viscosity ղ and the viscoelastic behavior (G’>>G’’) was evaluated as a function of time.

##### (iv) Assessment of sealing and mechanical properties of adhesive hydrogel in *ex vivo* porcine and rabbit eyes

The sealing and mechanical performance of the adhesive hydrogel was evaluated *ex vivo* in enucleated mature rabbit and porcine eyes using burst pressure measurements by using a modified ASTM standard, F2392-04. The porcine and rabbit eyeballs with extraocular tissues and intact optic nerve were custom ordered from Pelfreeze Biologics (Rogers, AR, USA) and shipped on ice in DPBS. All the burst pressure assessments were performed within 72 hours of arrival of the eyes using previously established procedure [59]. Briefly, the porcine and rabbit eyeballs were fixed on a custom-made holder under a Unitron Z650HR dissecting microscope (Feasterville, PA, USA). All the eyes were cannulated below the ora-serrata area into the vitreous cavity using a 27G needle connected with an IV tubing system which is connected to a syringe pump (NE1000, New Era Pump Systems Inc., Farmingdale, NY, USA) and a digital manometer (Digimano-1000, Netech, Farmingdale, NY, USA). The syringe and the tubing system up to the needle was filled with phosphate buffered saline. Five different wound architectures that mimic many aspects of ocular traumatic injuries were performed. The different incisions are (i) linear 5-6 mm full thickness central corneal incision, (ii) stellate 3.5 mm (3 pronged, each prong 3.5 mm) full thickness incision, (iii) circular (with central tissue) 4.5 mm full thickness incision, (iv) corneal full thickness perforation 3.0 mm and (v) 6 mm horizontal and 4mm vertical corneo-scleral incision. After the incision was performed, the cornea was gently pressed to remove aqueous humor and create anterior chamber hypotony to mimic some aspects of the warfare injury. The anterior chamber was gently filled with the injectable hydrogel via the incision using the cannula appropriately to recreate the chamber. Following recreation of the chamber, the adhesive pre-hydrogel solution was applied carefully between the tissue incision crevices and over the cornea covering the incision forming a layer extending over the healthy part of the tissue. The solution was photo crosslinked by illuminating the blue UV light for 90 seconds. The intraocular pressure was gradually raised by infusing saline (10mL/Hr) using the syringe pump. The burst pressure was noted from the manometer at the point when the infused fluid leaks from the incision point. The peelability of the adhesive hydrogel from the corneal surface was assessed qualitatively by a corneal surgeon by lifting and peeling the adhesive hydrogel layer using the surgical forceps. The results were documented using video and photographs.

##### (v) Assessment of wound closure and adhesive strength of the adhesive hydrogel

The wound closure capability of OcuPair adhesive hydrogel was evaluated by using a modified ASTM standard test F2458-05, Briefly, the corneal strips (1cm X 3 cm) including the cornea, and the sclera were prepared from freshly enucleated porcine eyeballs. A full thickness incision at the middle of the corneal strip. Approximately 25μL of the OcuPair pre-polymer adhesive hydrogel solution was applied between the two corneal strips (gap 1.2 mm) and over the corneal tissue (2 mm) and crosslinked using UV light for 90s on both sides. Parameters such as maximum load at break, time at break, and adhesion strength were evaluated using 5966 Dual Column Tabletop Testing System (Instron, Norwood, MA). The samples were clamped vertically and force from a 5-newton (N) load cell was applied at 0.05 N/min to stretch the sample until breaking. Lap shear strength was evaluated by using a modified lap shear test based on ASTM standard, F2255-05. The corneal strips were prepared (1cm X 4cm). A fill thickness incision was created using a razor blade. The two strips were adhered to each other with an overlapping distance of 0.5 cm facing the epithelial surface on both sides. The adhesive strength of the OcuPair adhesive hydrogel was measured at the detachment point.

#### 4.4.3. Stability analyses of the components of adhesive hydrogel (D-MA and HA-MA)

##### Stability study protocol

The stability analyses of D-MA and HA-MA samples were conducted over three months at three different temperatures: 4°C, 25°C, and 40°C. A concentration of 300 mg/mL and 190 mg/mL of D-MA and HA-MA respectively were prepared in a 10 mM phosphate buffer solution, aligning with the desired final concentration of the drug product. The samples were stored in amber-colored vials under argon atmosphere at the desired temperatures. At timepoints (1 week, 1-month, 2-months and 3-months), the samples were assessed using HPLC, pH measurements, osmolality, and ^1^H NMR.

##### HPLC methods for characterizing the D-MA & HA-MA conjugates

The purity of D-MA was determined using high-performance liquid chromatography (HPLC). The HPLC system (Waters Corporation, Milford, MA) was equipped with a 1525 binary pump, an In-Line degasser AF, a 717 plus autosampler, and a 2998 photodiode array (PDA) detector, managed by Waters Empower software. For analysis of D-MA conjugates, a Symmetry 300 C18 column (5 µm, 4.6 x 250 mm). The HPLC chromatograms starting dendrimer and D-MA conjugates were monitored at 210 (G4-OH) using the PDA detector. Mobile phase/solvent A: 0.1% TFA and 5% acetonitrile (ACN) in water, mobile phase/solvent B: 0.1% TFA in ACN. A gradient flow was used with initial condition of 100:0 (H2O/ACN) gradually changing the ratios to 50:50 (H2O/ACN) over 20 minutes, followed by 90:10 (H2O/ACN) at 30 minutes and the gradient was returned to initial conditions to 100:0 (H2O/CAN) at 45 minutes, with a constant flow rate of 1 mL/min.

For HA-MA conjugates, we have developed a robust HPLC method utilizing a size exclusion column (XBridge BEH200Å SEC 3.5 μm, 7.8 x 150 mm). For the mobile phase, we selected a 0.05N phosphate buffer, optimizing the separation process with a method runtime of 30 minutes. Monitoring the hyaluronic acid chromatogram at a wavelength of 210 nm, we observed the HA-MA eluting as a broad peak at 2.9 minutes (**Figure S4, Supporting information**). Each sample concentration was run in a triplicate to demonstrate its suitability. The line equation and the correlation coefficient (R²) were determined. An R² value above 0.98 was observed across the concentration range, demonstrating a strong linear relationship between the analyte concentration and the area under the curve (**Figure S10, Supporting information**).

### 4.5. *In vivo* assessment of OcuPair in a rabbit model of traumatic corneal injury

#### 4.5.1. Animals

All procedures involving animals conformed to the Association for Research in Vision and Ophthalmology Statement for the use of Animals in Ophthalmic and Vision Research and the study procedures were approved by the Johns Hopkins University Animal Care and Use Committee (protocol# RB17M53) and army ACURO (#USAMMDA-FY19-001). Fifty-six adult New-Zealand rabbits (Age 4-6 months, both males and females) were obtained from Harlan/Envigo Laboratories, Inc (IN, USA). All animals were housed at controlled temperature (16°C to 21°C), humidity (45% to 65%), standard 12/12 light/dark cycle and were fed with standard high fiber diet with hay and water.

#### 4.5.2. Surgical procedures

Survival surgeries on rabbits were performed under aseptic conditions. The right was used to create and treat the corneal wounds using OcuPair and Histoacryl whereas the left eye serves as control. All procedures were performed under general anesthesia (ketamine 50mg/kg (Bio-niche Pharma, Lake Forest, IL, USA) and (xylazine 5mg/kg (Phoenix Pharmaceuticals, St. Joseph, MO, USA) and topical anesthesia proparacaine 0.5% (Sandoz, Holzkirchen, Germany). Five different wounds/incisions were created on the cornea to mimic battle combat corneal injuries namely (i) linear full thickness gaping wound (5-6 mm), (ii) stellate (3.5mm) irregular laceration, (iii) a circular wound or PKP created using a 4.5 mm trephine in the central cornea, (iv) corneal-scleral (6 mm horizontal and 4mm vertical), and (v) corneal perforation (3mm). All incisions except circular PKP and corneal perforations were performed 24 hrs prior to the sealing to mimic battlefield injury situations. Also, before performing the incision the corneal surface and surrounding tissues were sterilized using 10% betadine and irrigated with sterile BSS. The corneal epithelium in the incision area was gently scrapped using Algerbrush II with 0.5mm burr tip (Surgeon’s Preference, MD, USA). All incisions were made using keratome surgical knife and cohan-vannas scissors. Damage to the lens/intraocular structures was carefully avoided. One dose of 10mg/kg acetazolamide (XGen Pharmaceuticals DJB, Inc, NY, USA) was administered via ear vein at the time of incision and other dose at 24hrs post incision (at the time of sealing with OcuPair). An anterior segment hypotony was created by removing the aqueous humor by incision and gently pushing the incision edge. The anterior chamber was filled with injectable gel (component 1 of OcuPair) to regain the anterior segment architecture. The incision was sealed by dispensing the adhesive hydrogel viscous pre-polymer solution (component 2 of OcuPair) via anterior chamber cannula, followed by blue UV light illumination for 90 seconds. The formation of the gel was assessed visually, and the sealing ensured by pressing the cornea and noticing if there is any leak of fluid from the aqueous chamber. Seidel test was performed to confirm the secure sealing. The performance of the adhesive hydrogel was evaluated against cyanoacrylate glue (Histoacryl, B.Braun, Germany). After stabilization, a contact ACCU Lens (16 mm dia and 6.0 bp, custom-made for rabbit corneas) (Masberrn Corp, CO, USA) was placed to avoid rabbits scratching the eyes. This was particularly useful for cyanoacrylate groups where that glue formed a rough surface over the cornea and caused irritation and to minimize the uncomfortable pain during blinking. Dexamethasone 0.1% (Bausch + Lomb) and moxifloxacin 0.5% (Aurobindo) were administered twice a day for all eyes with procedures, and 0.25% buprenorphine were administered subcutaneously every 12 hours for 48 hours as per the JHU protocol requirement. The eyes were inspected for chamber formation, iris location, corneal clarity, presence and integrity of glue, changes in IOP, signs of leakage from site of incision, signs of corneal infections and inflammation at day 3 and day 5 post surgery.

#### 4.5.3. Slit lamp microscopy

Slit lamp microscopy and photography was performed under general anesthesia using a Haag-Streit slit lamp system. Photographs were also taken at the time of examination. Clinical parameters such as anterior chamber formation, corneal opacity, corneal epithelial edema, IOP measurements, conjunctival chemosis and Flare were evaluated and scored according to the scoring system represented in **table S5, supporting information**. In addition to clinical parameters, OcuPair adhesive hydrogel’s performance was also evaluated in terms of hydrogel integrity, its adherence to the corneal tissue, visibility of intraocular contents vis hydrogel bandage and integrity of the hydrogel bandage. To access if there is any leakage from the site of incision, we performed Seidel leakage test my staining the cornea using fluoresceine eye drops (ALTAFLUOR BENOX) and cobalt blue slit lamp photography.

#### 4.5.4. Intraocular pressure measurements

The intraocular pressure was measured using a handheld tonometer Icare® TA01i (iLab tonometer, iCare, Finland). Rabbit settings were chosen, and the tonometer was used according to the manufacturer’s instructions.

#### 4.5.5. Anterior segment – Optical coherence tomography (AS-OCT)

The central corneal thickness was evaluated using anterior segment optical coherence tomography (Bioptigen, NC, USA) equipped with 10mm anterior telescopic lens. The central corneal thickness (CCT) was measured using Bioptigen software.

#### 4.5.6. Toxicity and biocompatibility evaluation of OcuPair adhesive hydrogel

##### (i) Histopathology evaluation

The rabbits were euthanized at POD3 and POD5 and the eyes were enucleated and fixed in 10% PFA fixative solution, using our lab’s established procedures to preserve the structure of the eye globe [38]. After fixation, the extra tissues were carefully trimmed on both sides of the incision area to obtain eye rings and were given to Johns Hopkins pathology core for paraffin embedding, sectioning, and H&E staining. Ten microns sections at the incision site (2-3 sections for each eye) were used for analysis. The sections were imaged under Nikon Eclipse Ni microscope for signs of changes in corneal tissue, corneal epithelium, stromal edema and signs of inflammation (macrophages and neutrophils accumulation).

##### (ii) Thirty-day corneal and ocular toxicity

Corneal and intraocular toxicity of OcuPair adhesive hydrogel and injectable hydrogel was evaluated in a GLP setting at Charles rivers (as part of army contract, in accordance with ISO 10993-6) using GLP engineering batches (Developed at GLP-CRO facility following the optimized synthesis protocol reported in this manuscript) in adult New-Zealand rabbits (36 rabbits (18 males & 18 females), not included in the JHU study). To evaluate the intraocular toxicity of injectable filler gel, ∼50µL and ∼40µL were injected into anterior and posterior chamber respectively. For adhesive hydrogel solution, ∼20µL was injected into the anterior chamber. For evaluation the toxicity of adhesive hydrogel in gel form, ∼50µL of the solution was injected into subconjunctival space followed by photo-crosslinking the injected solution using UV light exposure for 4 minutes to form a hydrogel under the subconjunctival space. Clinical parameters such as intraocular uveitis, changes in intraocular pressure, corneal toxicity and edema were evaluated for a period of 30 days.

##### (iii) Biocompatibility evaluation

Long term biocompatibility and corneal toxicity was evaluated in adult New-Zealand rabbits (n=3) by performing lamellar keratoplasty in the central cornea. A 4.5 mm partial keratoplasty with ∼40-50 % of stroma was removed to form a well. The space was filled with ∼75μL of OcuPair adhesive pre-polymer solution and photo-crosslinked with blue UV light for 90 seconds resulting in a transparent hydrogel. The animals were monitored for signs of cornel toxicity, corneal re-epithelization, stromal integration, and biocompatibility were assessed using slit lamp microscopy, Seidel test and AS-OCT as previously mentioned on POD 10, 30, and 4 months.

## 5. Statistical analyses

All data are presented as means ± SEM. The analyses were conducted in Excel 2013 and GraphPad Prism (version 6; La Jolla, CA). Student’s *t*-tests for different single groups and One-way ANOVA were performed for multiple-group comparisons, with Tukey’s post hoc test: **P < 0.05, and ***P < 0.001.

## 7. Supplementary Information

**The supplementary information PDF includes the following:**

Fig S1: 1H NMR spectra of different D-MA pilot scale batches.

Fig S2: HPLC chromatogram of D-MA.

Fig S3: 1H NMR spectra of different HA-MA pilot scale batches.

Fig S4: HPLC chromatogram of HA-MA.

Fig S5: Viscosity measurements of OcuPair injectable hydrogel and adhesive hydrogel solutions.

Table S1: Summary of findings in the process of optimizing the OcuPair adhesive hydrogel formulation.

Fig S6: Stability evaluation of D-MA solution using HPLC. Fig S7: Stability evaluation of HA-MA solution using HPLC.

Table S2: Summary of stability evaluation if D-MA and HA-MA up to 3-month timepoint at different storage temperatures.

Table S3: Summary of *ex vivo* eye burst pressure measurements in rabbit and porcine eyeballs.

Fig S8: Evaluation of effect of D-MA degradation during stability studies on burst pressure measurements in ex vivo porcine eyes.

Fig S9: Photographs of OcuPair injectable hydrogel and adhesive hydrogel pre-formulation loaded into final delivery devices as part of OcuPair kit.

Fig S10: Calibration graph for HA-MA using HPLC analysis. Table S4: Clinical parameters and their grading score system

## Other supplementary materials include the following

Video 1: Manipulation of the rabbit eye with linear incision sealed with OcuPair adhesive hydrogel.

Video 2: Manipulation of the rabbit eye with stellate incision sealed with OcuPair adhesive hydrogel.

Video 3: Peeling of OcuPair adhesive hydrogel from the porcine eyes using surgical forceps.

## Supporting information

Video1: Linear wound with OcuPair

Video2: Stellate wound with OcuPair

Video3: OcuPair peeling

## 8. Acknowledgements & Funding

The authors would like to acknowledge Wilmer-Woods animal facility staff for animal housing and procedures. **Funding:** This work was supported by is funded under contract W81XWH-18-C-0180 (R.M.K., and S.P.K.,) with U.S. Army Medical Material Development Activity (USAMMDA), Temporary Corneal Repair (TCR) division, Foundation for Fighting Blindness, Arnall Patz Distinguished Professorship funds, and Dr. Walter Stark’s gift funds.

## 9. Author contributions

R.M.K., S.P.K., and S.C.Y., conceived the idea and designed the research study. R.M.K. and S.P.K. secured funding from USAMMDA-TCR and supervised the research. S.P.K., and R.S., performed the synthesis, characterization and pilot-scale process development. S.P.K optimized the formulation and performed the rheology and mechanical testing experiments. S.P.K., and H.L., carried out the *ex vivo* and *in vivo* experiments. R.S., and S.P.K collaborated with CMO and facilitated scale-up of OcuPair components. S.A., assisted with synthesis and purification of OcuPair. S.P.K, and J.L.C., collaborated with CRO and supervised the GLP toxicity study. S.P.K., R.S., R.M.K., and S.C.Y wrote the manuscript, and all the others edited the manuscript.

## 10. Competing interests/Conflict of interests

The authors have awarded and pending patents relating to the hydroxyl dendrimer platform (R.M.K., and S.P.K.) and dendrimer-bioadhesive polymer hydrogel for ocular applications (R.M.K., S.P.K., and S.C.Y.). R.M.K., and his wife (Sujatha Kannan) are co-founders and have financial interests in Ashvattha Therapeutics; a start-up translating dendrimer-drug delivery platform. J.L.C. is the president and CEO of Ashvattha Therapeutic Inc. R.S., and S.A. were full-time employee of Ashvattha Therapeutic Inc. All other authors declare no competing or conflict of interests.

## 11. Data and materials availability

All data needed to evaluate the conclusions in the paper are present in the paper and/or the Supplementary Materials. The data that support the findings of this study are available from the corresponding authors upon reasonable request.

