## Supporting or supplemental information for "OcuPair, a novel photo-crosslinkable dendrimer-hyaluronic acid hydrogel bandage/bioadhesive for corneal injuries and temporary corneal repair"

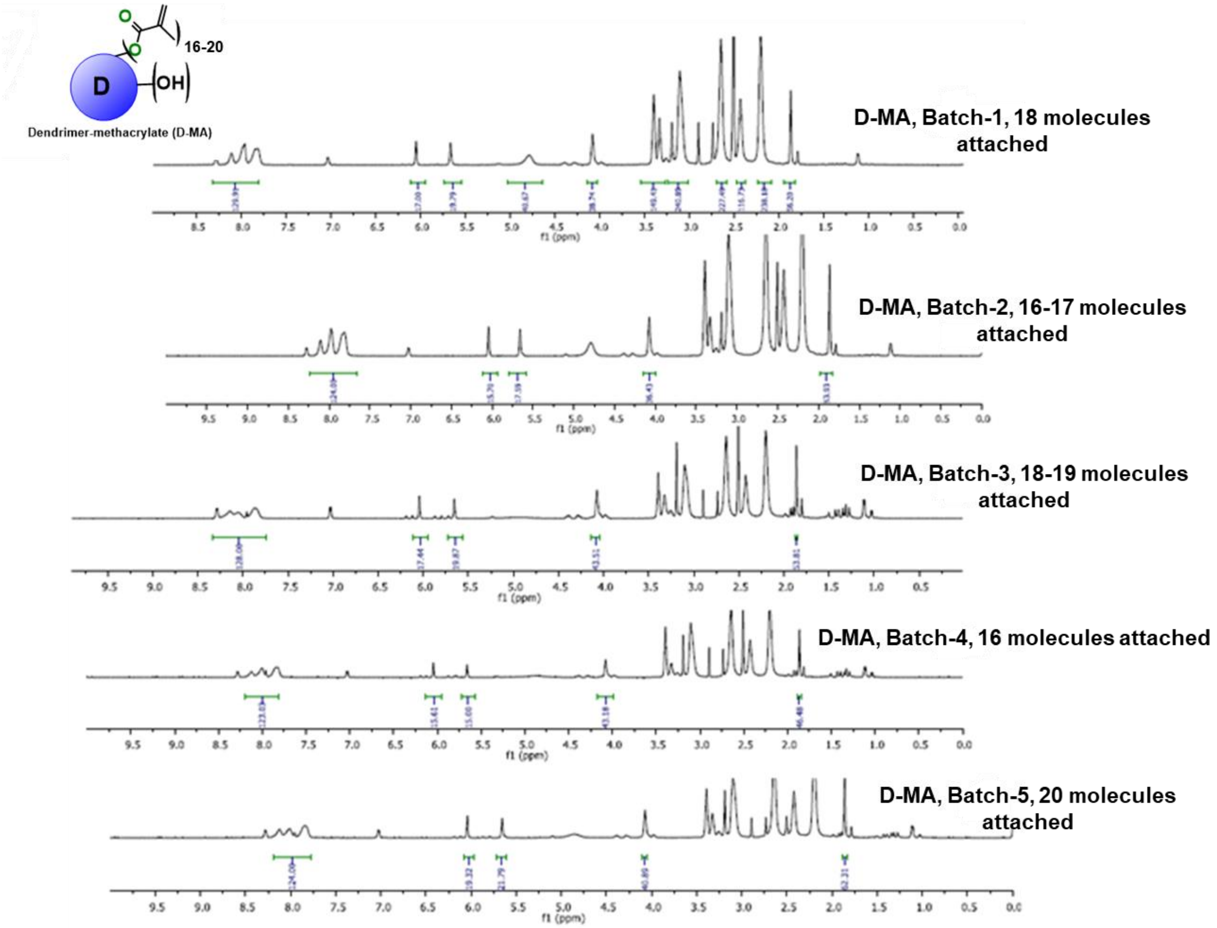


**Figure S1:** [1]H NMR spectra of different batches of D-MA synthesized using the optimized protocol reported demonstrating consistent methacrylate loading.

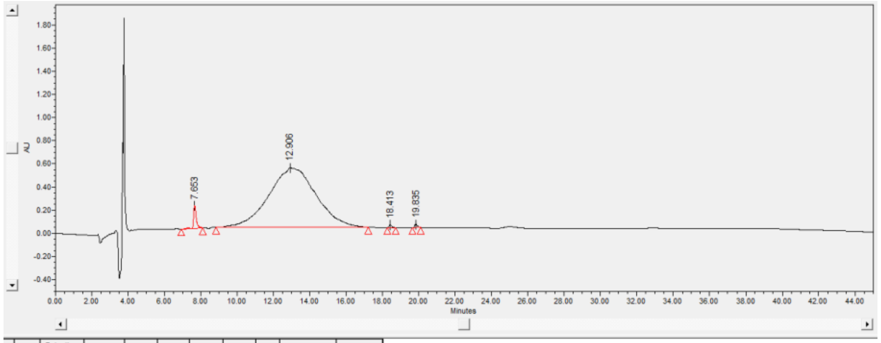


| | Name | Retention Time (min) | Area (µV*sec) | % Area | Height (µV) | Int Type | Amount | Units | Peak Type | Peak Codes |
|---|---|---|---|---|---|---|---|---|---|---|
| 1 | | 7.653 | 2020779 | 2.13 | 195824 | bb | | | Unknown | |
| 2 | | 12.906 | 92438673 | 97.27 | 514835 | bb | | | Unknown | |
| 3 | | 18.413 | 256525 | 0.27 | 23950 | bb | | | Unknown | |
| 4 | | 19.835 | 317881 | 0.33 | 29672 | bb | | | Unknown | |

**Figure S2:** HPLC chromatogram of synthesized D-MA conjugate demonstrating a purity >95% eluting at 12.9 min.

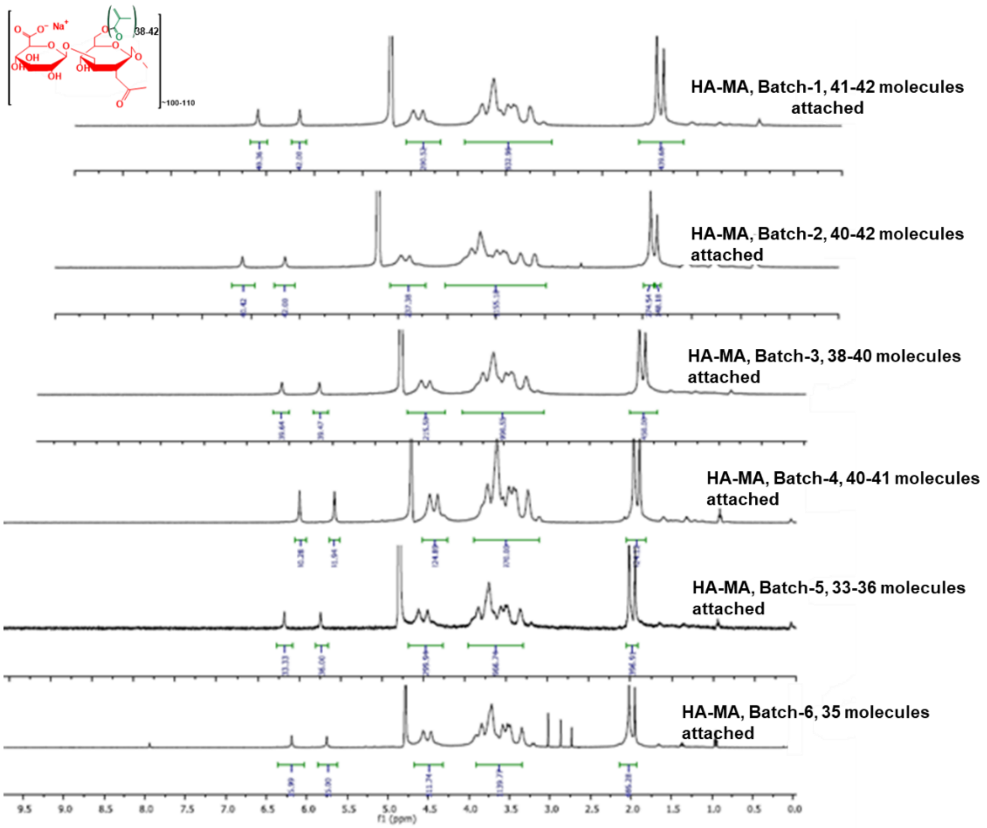


**Figure S3:** [1]H NMR spectra of different batches of HA-MA synthesized using the optimized protocol reported demonstrating consistent methacrylate loading.

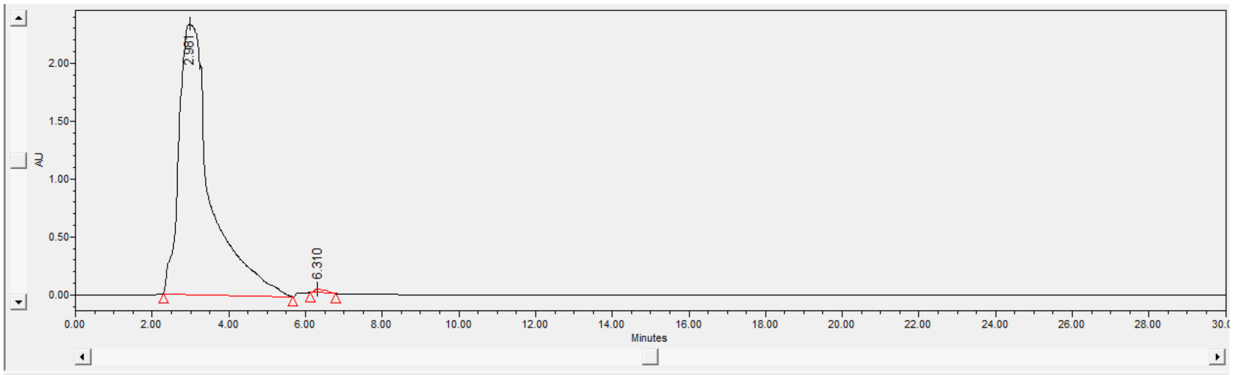


| | Name | Retention Time (min) | Area (µV*sec) | % Area | Height (µV) | Int Type | Amount | Units | Peak Type | Peak Codes |
|---|---|---|---|---|---|---|---|---|---|---|
| 1 | | 2.981 | 134669709 | 99.52 | 2330408 | bb | | | Unknown | |
| 2 | | 6.310 | 647116 | 0.48 | 33283 | bb | | | Unknown | I08 |

**Figure S4:** HPLC chromatogram of synthesized HA-MA conjugate demonstrating purity >97% eluting at 2.98 min.

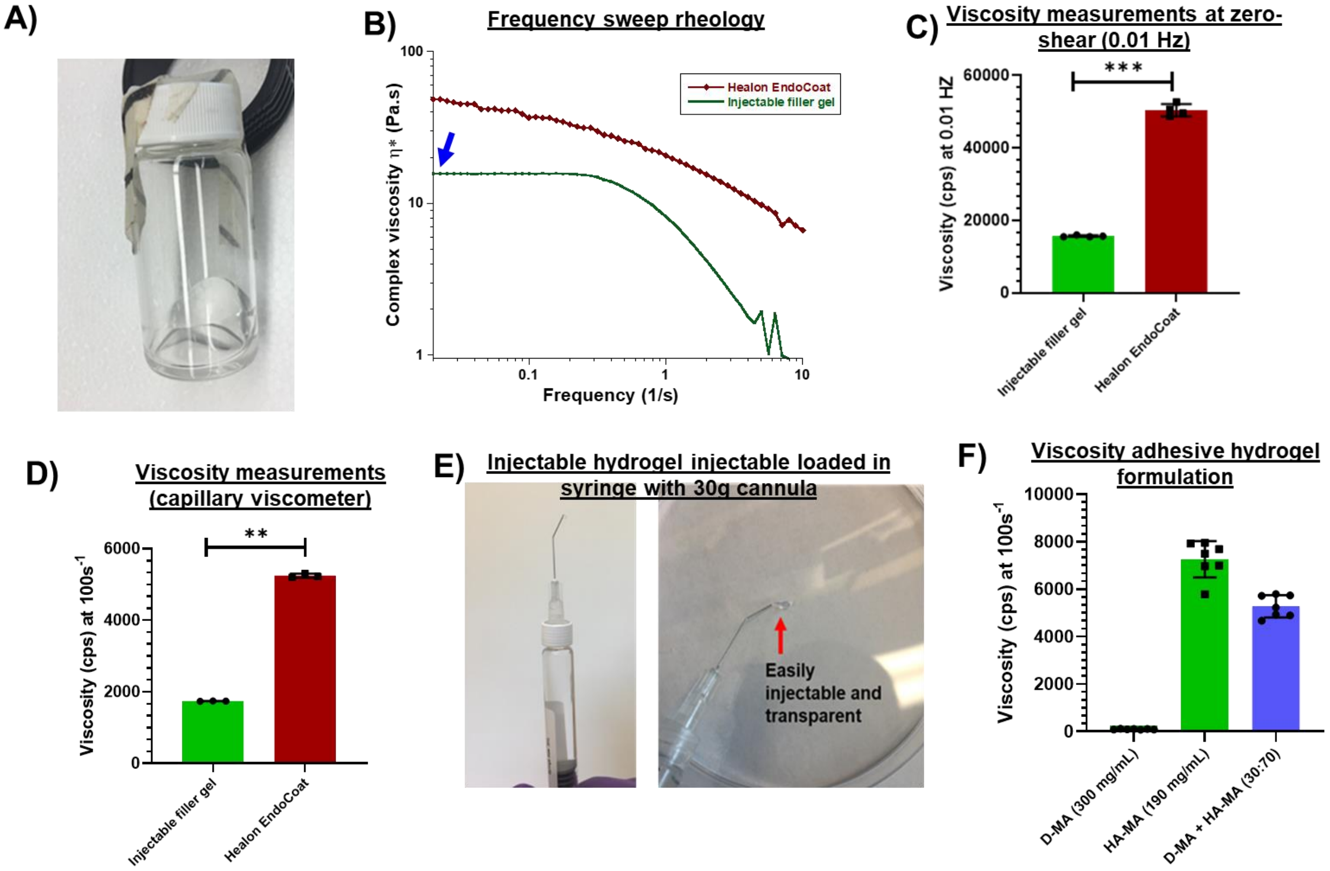


**Figure S5: Viscosity measurements of OcuPair injectable hydrogel and adhesive hydrogel formulation. A)** Image of injectable hydrogel after sterilization demonstrating that the solution is transparent and viscous. **B)** Dynamic frequency sweep rheology of injectable hydrogel and using parallel plate rheometer demonstrating shear thinning properties at low viscosity compare the Healon Endocoat. **C)** Complex viscosity measurements at zero shear frequency of 0.01 Hz for injectable hydrogel and compared with Healon Endocoat. **D)** Viscosity of Injectable hydrogel formulation at constant frequency of 100 $s^{-1}$ using capillary viscometer. **E)** Image of injectable hydrogel loaded into a glass syringe fitted with 30G anterior chamber cannula demonstrating that the injectable hydrogel can be easily extruded through narrow gauge needle. **F)** Viscosity measurements of D-MA (300 mg/mL), HA-MA (190 mg/mL) and adhesive hydrogel formulation solutions at constant frequency of 100 $s^{-1}$.

**Table S1:** Summary of findings in the process of optimizing the OcuPair adhesive hydrogel formulation.

| D-MA: HA-MA ratio in the formulation | Viscosity (cps) | Gelation time (seconds) | Gel characteristics and appearance | Residence time after applying on the cornea | Burst pressure in rabbit eyes (mmHg) for linear incision | Image |
|---|---|---|---|---|---|---|
| 100% D-MA (100:0) | ~96.8 ± 6.6 (n=7) | ~10-20 s | Forms thin and brittle gels and the gel breaks when twisted using forceps | The solution flows away or flows into the corneal incision immediately applying | NA | 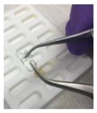 |
| 100% HA-MA (0:100) | ~7306.5 ± 336.5 (n=12) | >200 s | Forms patchy weak gels | The solution stays on the corneal incision for more than 20 s after application | NA | 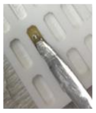 |
| 70:30 | ~4142.3 ± 185 (n=6) | ~30-45 s | Forms brittle gels and gels breaks when handled | The solution stays on corneal surface for ~10-15s | ~42.8 ± 7.2 mmHg The hydrogel layer peels from corneal surface quickly at burst pressure | 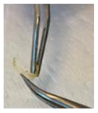 |
| 50:50 | ~5068 ± 87 (n=6) | ~30-45 s | Forms flexible gels but some gels breaks when handled | The solution stays on corneal surface for ~15-20 s after application | ~58.4 ± 6.8 mmHg The hydrogel layer peels from corneal surface at burst pressure | 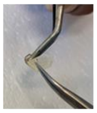 |
| 30:70 | ~5205.8 ± 190 (n=6) | ~60-80 s | Forms flexible and transparent and sticky gels and can be easily handled using instruments | The solution stays on corneal surface for ~15-20 s upon application | ~78.4 ± 2.6 mmHg. The hydrogel bandage peels from the corneal surface at burst pressure but withstands ~60mmHg for 3-5 mins | 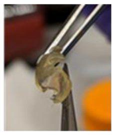 |

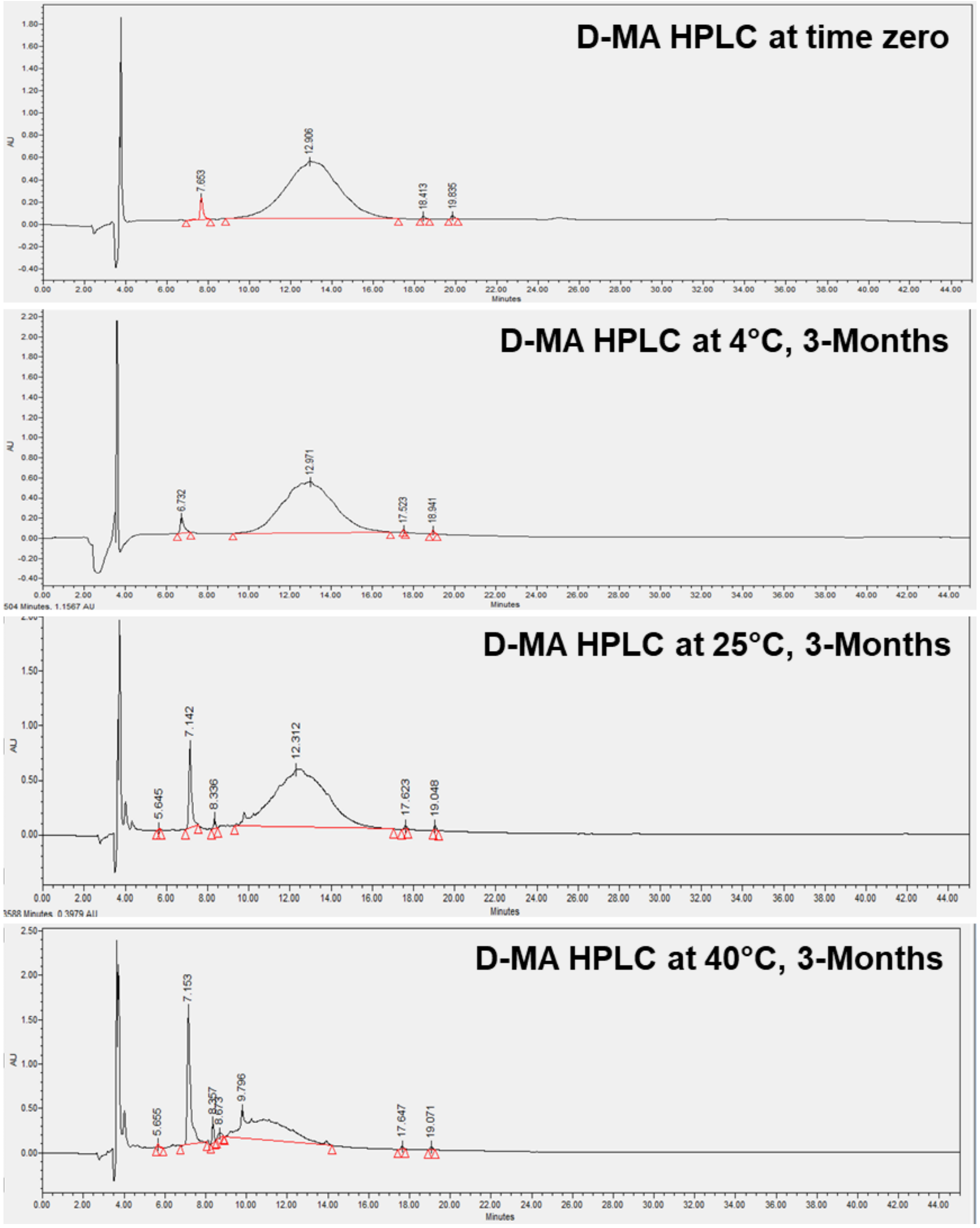


**Figure S6: Stability evaluation of D-MA solution using HPLC.** HPLC chromatograms of D-MA (300 mg/mL) solution in phosphate buffer (pH 6) stored at 4ºC, 25ºC, and 40ºC at 3-month time point compared with time zero.

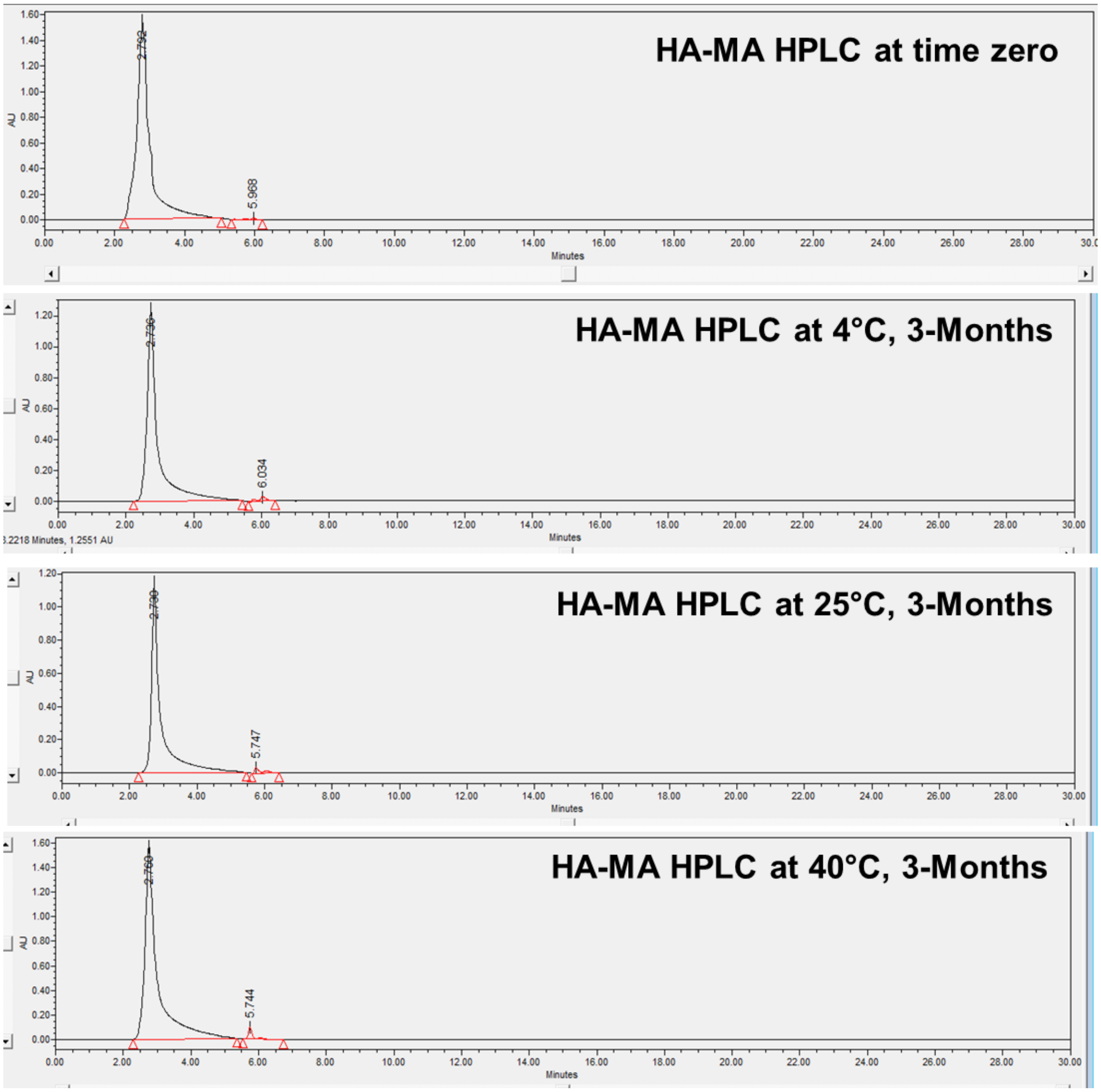


**Figure S7: Stability evaluation of HA-MA solution using HPLC.** HPLC chromatograms of HA-MA (190 mg/mL) solution in phosphate buffer (pH 6) stored at 4ºC, 25ºC, and 40ºC at 3-month time point compared with time zero.

| Summary of stability evaluation of Dendrimer-Methacrylate (D-MA) solution (300mg/mL) | | | | | | | | | |
|---|---|---|---|---|---|---|---|---|---|
| Time points/Temperature | 4°C | | | 25°C | | | 40°C | | |
| Parameters | HPLC purity (%) | pH | Osmolality (moSm/kg) | HPLC purity (%) | pH | Osmolality (moSm/kg) | HPLC purity (%) | pH | Osmolality (moSm/kg) |
| Time Zero | 97.3 | 7.59 | 201 | | | | | | |
| 1 week | 97.5 | 7.37 | 208 | 96.8 | 7.42 | 237 | 94.8 | 7.40 | 300 |
| 1 month | 97.6 | 7.39 | 200 | 95.6 | 7.48 | 236 | 78.7 | 7.31 | 382 |
| 2 months | 97.6 | 7.37 | 215 | 92.2 | 7.35 | 255 | 74.3 | 7.63 | 396 |
| 3 months | 97.3 | 7.31 | 205 | 92.3 | 7.47 | 249 | 65.4 | 7.23 | 356 |
| Summary of stability evaluation of Hyaluronic acid-Methacrylate (HA-MA) solution (190mg/mL) | | | | | | | | | |
| Time points/Temperature | 4°C | | | 25°C | | | 40°C | | |
| Parameters | HPLC purity (%) | pH | Osmolality (moSm/kg) | HPLC purity (%) | pH | Osmolality (moSm/kg) | HPLC purity (%) | pH | Osmolality (moSm/kg) |
| Time Zero | 99.5 | 7.61 | | | | | | | |
| 1 week | 99.3 | 7.42 | | 99.6 | 7.34 | | 99.0 | 7.59 | |
| 1 month | 99.2 | 7.37 | | 98.6 | 7.61 | | 98.5 | 7.43 | |
| 2 months | 98.4 | 7.55 | | 98.9 | 7.54 | | 98.2 | 7.48 | |
| 3 months | 98.3 | 7.39 | | 98.0 | 7.29 | | 98.0 | 7.67 | |

**Table S2:** Summary of stability evaluation if D-MA and HA-MA up to 3-month timepoint at different storage temperatures.

**Table S3:** Summary of *ex vivo* eye burst pressure measurements in rabbit and porcine eyeballs.

| **Summary of ex vivo eye burst pressure measurements** | | |
|---|---|---|
| **Incision Type** | **Porcine Eyes (mature)** | **Rabbit eyes (mature)** |
| **Linear full thickness (5-6mm)** | 101.5 ± 6.5 mmHg (n=18) | 78.4 ± 2.6 mmHg (n=16) |
| **Stellate (three pronged) (3.5 mm each prong)** | 92.6 ± 5.8 mmHg (n=18) | 74.7± 5.4 mmHg (n=16) |
| **Circular incision with tissue present (4.5mm)** | 84.2 ± 9.3 mmHg (n=18) | 58.4 ± 3.0 mmHg (n=12) |
| **Corneal perforations (3mm dia)** | 115 ± 5.3 mmHg (n=16) | 86.2 ± 7.2 mmHg (n=13) |
| **Corneal-scleral incisions 6mm horizontal and 4mm vertical** | >120 mmHg (n=18) | >100 mmHg (n=15) |

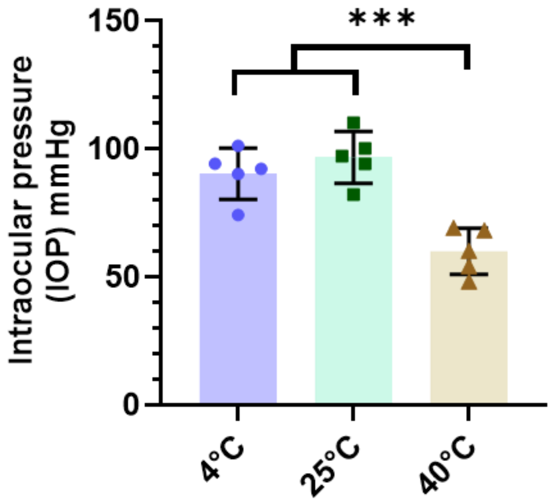


**Figure S8: Evaluation of effect of D-MA degradation during stability studies on burst pressure measurements in ex vivo porcine eyes.** The adhesive hydrogel formulation was prepared by using the D-MA solution (300mg/mL) stored for 3 months at 4°C, 25°C and 40°C.

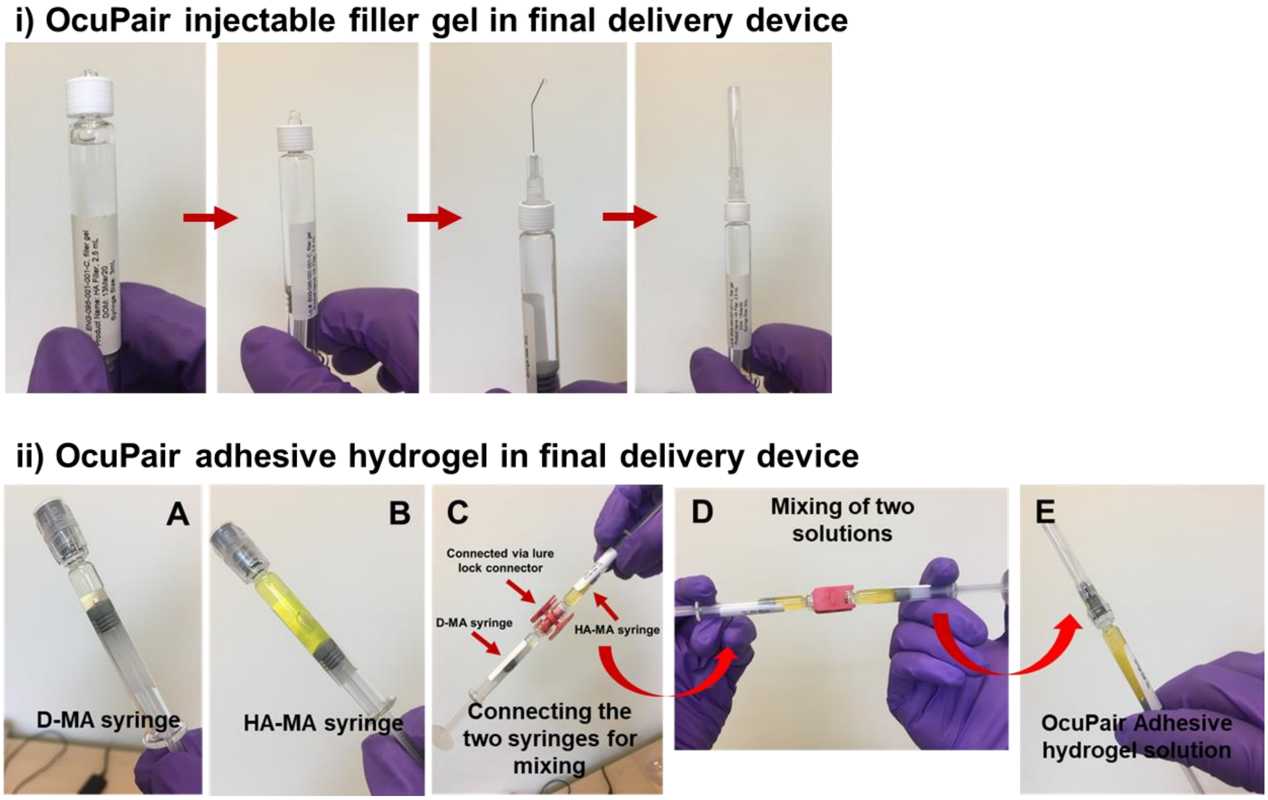


**Figure S9: OcuPair injectable hydrogel and adhesive hydrogel pre-formulation loaded into final delivery devices as part of OcuPair kit. (i)** OcuPair injectable hydrogel (sterilized) loaded into 3mL glass syringes with lure-lok connector under sterile conditions. A 30G anterior chamber cannula cane connected to the syringe and the injectable hydrogel can be easily injected into the intraocular cavity. **(ii)** The individual components of the adhesive hydrogel D-MA (300 mg/mL) and HA-MA (190mg/mL) with 0.05% fluoresceine (hence the yellow color) were loaded in to separate 1 mL glass syringes with luer-lok caps aseptically. The two solutions are mixed by connecting the two syringes using a lure-lok connector and all the contents are pushed into one syringe and fitted with a 27G cannula for applying over the corneal wound. All the components are provided as part of OcuPair kit.

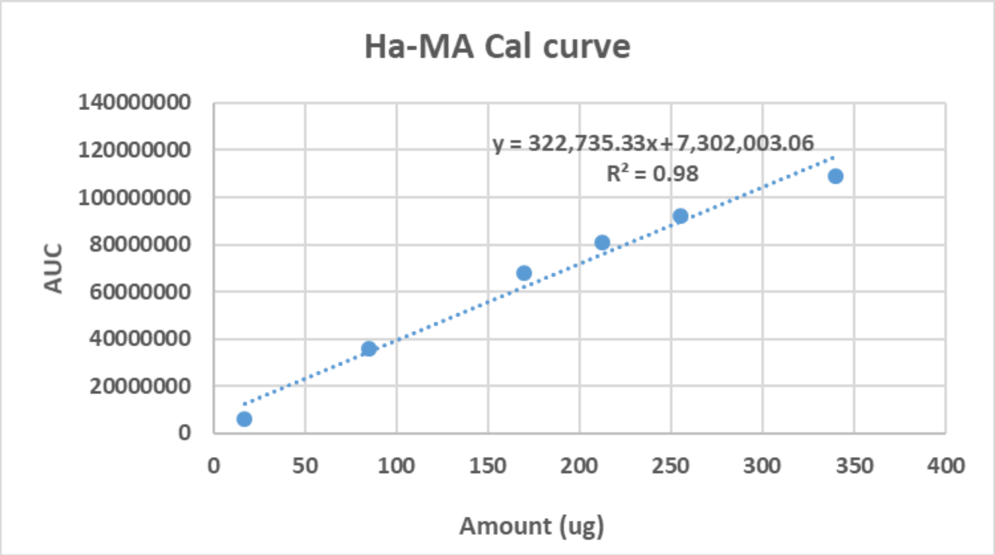


**Figure S10:** Calibration graph for HA-MA using HPLC analysis.

| Clinical Parameters | Scores |
|---|---|
| **Anterior chamber formation** | 0 = no visible chamber, 1 = shallow chamber (30-50% depth), 2 = mild shallow chamber (50-80% depth), 3 = full chamber (>80% depth) |
| **Corneal opacity** | 0 = no opacity and completely clear, 1 = slight haze with iris and lens visible, 2 = moderately opaque, iris and lens detectable, 3 = severely opaque with iris and lens hardly visible. |
| **Conjunctival chemosis** | 0 = no chemosis, 1+ = mild chemosis, 2+ = moderate chemosis, 3+ = severe chemosis |
| **Corneal epithelial edema** | 0 = no edema, 1+ = mild edema, 2+ = moderate edema, 3+ = severe edema |
| **FLARE (corneal and anterior chamber inflammation)** | 0 = no flare, 1+ = faint flare, 2+ = moderate flare (iris and lens details are clear), 3+ = marked flare (iris and lens details are hazy), 4+ = intense flare (with fibrin exudate) |

**Table S4:** Clinical parameters and their grading score system

**Legends for videos**

**Video 1:** Rabbit eyeball with full-thickness linear (6-8mm) wound sealed using OcuPair adhesive hydrogel.

**Video 2:** Rabbit eyeball with full-thickness stellate wound sealed using OcuPair adhesive hydrogel.

**Video 3**: Peeling of adhesive hydrogel bandage from the corneal surface using surgical forceps.
